# Human Pancreatic α-Cell Heterogeneity and Trajectory Inference Analysis Using Integrated Single Cell- and Single Nucleus-RNA Sequencing Platforms

**DOI:** 10.1101/2023.11.19.567715

**Authors:** Randy B. Kang, Jungeun Lee, Miguel Varela, Yansui Li, Carolina Rosselot, Tuo Zhang, Esra Karakose, Andrew F. Stewart, Donald K. Scott, Adolfo Garcia-Ocana, Geming Lu

**Affiliations:** Department of Molecular and Cellular Endocrinology, Arthur Riggs Diabetes and Metabolic Research Institute, Beckman Research Institute of City of Hope, Duarte, CA, 91010; Diabetes Obesity and Metabolism Institute, and Division of Endocrinology, Diabetes and Bone Diseases, Icahn School of Medicine at Mount Sinai, New York, NY, 10029; Genomics Resources Core Facility, Weill Cornell Medicine, New York, NY, 10065

## Abstract

Prior studies have shown that pancreatic α-cells can transdifferentiate into β-cells, and that β-cells de-differentiate and are prone to acquire an α-cell phenotype in type 2 diabetes (T2D). However, the specific human α-cell and β-cell subtypes that are involved in α-to-β-cell and β-to-α-cell transitions are unknown. Here, we have integrated single cell RNA sequencing (scRNA-seq) and single nucleus RNA-seq (snRNA-seq) of isolated human islets and human islet grafts and provide additional insight into α-β cell fate switching. Using this approach, we make seven novel observations. 1) There are five different *GCG*-expressing human α-cell subclusters [α1, α2, α-β-transition 1 (AB-Tr1), α-β-transition 2 (AB-Tr2), and α-β (AB) cluster] with different transcriptome profiles in human islets from non-diabetic donors. 2) The AB subcluster displays multihormonal gene expression, inferred mostly from snRNA-seq data suggesting identification by pre-mRNA expression. 3) The α1, α2, AB-Tr1, and AB-Tr2 subclusters are enriched in genes specific for α-cell function while AB cells are enriched in genes related to pancreatic progenitor and β-cell pathways; 4) Trajectory inference analysis of extracted α- and β-cell clusters and RNA velocity/PAGA analysis suggests a bifurcate transition potential for AB towards both α- and β-cells. 5) Gene commonality analysis identifies *ZNF385D, TRPM3, CASR, MEG3* and *HDAC9* as signature for trajectories moving towards β-cells and *SMOC1, PLCE1, PAPPA2, ZNF331, ALDH1A1, SLC30A8, BTG2, TM4SF4, NR4A1* and *PSCK2* as signature for trajectories moving towards α-cells. 6) Remarkably, in contrast to the events *in vitro*, the AB subcluster is not identified *in vivo* in human islet grafts and trajectory inference analysis suggests only unidirectional transition from α-to-β-cells *in vivo*. 7) Analysis of scRNA-seq datasets from adult human T2D donor islets reveals a clear unidirectional transition from β-to-α-cells compatible with dedifferentiation or conversion into α-cells. Collectively, these studies show that snRNA-seq and scRNA-seq can be leveraged to identify transitions in the transcriptional status among human islet endocrine cell subpopulations *in vitro*, *in vivo*, in non-diabetes and in T2D. They reveal the potential gene signatures for common trajectories involved in interconversion between α- and β-cells and highlight the utility and power of studying single nuclear transcriptomes of human islets *in vivo*. Most importantly, they illustrate the importance of studying human islets in their natural *in vivo* setting.

## INTRODUCTION

Glucagon-producing pancreatic α-cells were discovered seventy-five years ago and are recognized as important physiological regulators of life-threatening hypoglycemia by counteracting the effects of insulin on glucose homeostasis (1,2). In diabetes, patients display postprandial hyperglucagonemia which exacerbates hyperglycemia (3–6). Despite these important observations, research on cellular heterogeneity in the human pancreatic islet has mainly focused on insulin-producing β-cells, and studies describing the range of α-cell subpopulations are scarce. Recently, however, this has changed and specific human α-cell subpopulations that participate in normal and abnormal glucose homeostasis have attracted attention. For example, recent evidence indicates that there is variation in the glucagon content among human α-cells (7) and that α-cell functional heterogeneity is linked to α-cell maturation in type 2 diabetes (T2D) (8). Furthermore, an increasing subpopulation of glucagon-like peptide-1 secreting α-cells occurs in T2D (9). Finally, elevated serum amino acids have been reported to induce a subpopulation of α-cells to initiate pancreatic neuroendocrine tumor formation (10). Collectively, these studies indicate that deeper analysis of α-cell heterogeneity and the mechanisms controlling their identity in health and disease is warranted.

Single-cell RNA sequencing (scRNA-seq) has revolutionized the identification and analysis of different cell types within heterogeneous populations. By algorithmically clustering the data, it has been possible to annotate cell types, and with the use of different clustering parameters such as Louvain resolution, it has been possible to investigate subpopulations with distinct transcriptomes in an unbiased manner (11). In the human islet, scRNA-seq has uncovered several α- and β-cell subtypes with different transcriptome profiles that can predict the maturity of these islet cell subtypes in basal and diabetes conditions (12–17). However, how these α- and β-cell subtypes can transcriptionally transition from one cell subtype to another remains understudied.

Loss of pancreatic β-cells leads to diabetes. α-cells are refractory to spontaneous conversion to β-cells, but a small percentage of α-cells are reprogrammed to β-cells with acute β-cell loss (18). β-cell replacement by reprogramming α-cells into β-cells may be a promising approach to diabetes therapy and some researchers have focused on uncovering the mechanisms controlling adult human α-cell identity (19–22). Human islet β-cell subpopulations with different dynamic transcriptome profiles have been recently identified (15–17,23,24). However, whether specific subpopulations of human α-cells and β-cells have the potential to transcriptionally evolve into any other subpopulation under basal conditions is understudied. Furthermore, since most human islet studies are performed on islets isolated from organ donors, events that occur *in vivo* in human islet grafts are unknown.

Against this background, we have shown that integration of scRNA-seq and single nucleus RNA-seq (snRNA-seq), and the use of intronic reads can be of great value for identifying β-cell subpopulations in human islets not only *in vitro* but also in human islet grafts *in vivo* (17). Here, using this scRNA-seq and snRNA-seq reference dataset, we identify five different *GCG*-expressing α-cell subtypes with different transcriptome profiles in human islets isolated from adult non-diabetic donors. We find that one of the subpopulations identified *in vitro*, α/β-cells (AB cells) is a multihormonal α-cell subpopulation that can potentially transition in a bifurcated manner into both mature α- and β-cells. However, AB cells are not present in human islet grafts *in vivo* and trajectory analysis shows that transitions are only unidirectional from α-cells to β-cells in this setting. Using trajectory analysis of the different α-and β-cell subpopulations, we identify *ZNF385D, TRPM3, CASR, MEG3* and *HDAC9* as common genes for the trajectories from α- to β-cells. Conversely, we find S*MOC1, PLCE1, PAPPA2, ZNF331, ALDH1A1, SLC30A8, BTG2, TM4SF4, NR4A1* and *PSCK2* as genes identifying cells on a trajectory from β- to α-cells. Further, trajectory analysis of human islet cells isolated from T2D donors shows a unidirectional transition from β-cells to α-cells. Expression of two α-cell common genes, *SMOC1* (sparc-related modular calcium-binding protein 1) and *ALDH1A1* (aldehyde dehydrogenase 1 isoform A1) are increased in T2D β-cells, and their expression negatively correlate with *INS* expression. Collectively, these studies highlight the usefulness of the integrated scRNA-seq and snRNA-seq reference to identify α-cell subpopulations *in vitro* and *in vivo*. It facilitates the analysis of potential transitions between α-cells and β-cells at a transcriptional level depending on the pathophysiological context. They also highlight *SMOC1* and *ALDH1A1* as potential dedifferentiation genes in β-cells.

## MATERIAL AND METHODS

### Human islet samples

Adult human pancreatic islets from eleven brain-dead donors were isolated by Prodo Laboratories (Aliso Viejo, CA) according to the standard procedure and used for these studies (**Additional file 1: Table S1**) (17,25). Briefly, islets were harvested from pancreata from deceased organ donors without any identifying information and with informed consent properly and legally secured, and Western Institutional Review Board (WIRB) approval. The average donor age was 45 ± 5, 73% were male donors and additional donor demographic information is included in **Additional file 1: Table S1**. Human pancreatic islets from three non-diabetic donors were used for the *in vitro* studies (detailed below) and human islets from eight donors were used in human islet transplant experiments in immunosuppressed mice (detailed below). Transcriptomics analysis of some of these samples has been previously published (17). In addition, we mined the raw FASTQ data from Human Pancreas Analysis Program (HPAP) of the Human Islet Research Network (HIRN), and we have performed analysis of 13 non-diabetic and 13 T2D scRNA-sequencing datasets through SFTP (secure file transfer protocol) (details below) (26).

### Human islet cells, human islet transplantation into RAG-1^−/−^ immunodeficient mice and nuclei processing

Human islets from three different donors (3000 IEQs/donor) (**Additional file 1: Table S1**) were used in these studies as previously reported (17). Briefly, islet cells were dissociated using pre-warmed Accutase (cat# 25–058-CL, Corning) and half of the cells were resuspended in binding buffer (cat# 130–090-101, Miltenyi Biotec) with dead cell removal beads, and applied onto the dead cell removal column (cat # 130–042-401, Miltenyi Biotec), which was attached to the MACS separator. Then the cell concentration was measured with the Countess-3 Automated Cell Counter (Thermo-Fisher). The other half of the cells was homogenized with a pestle, and their nuclei were isolated with the Minute™ single nucleus isolation kit for tissue/cells (Cat# SN-047, Invent Biotechnologies, INC) and the nuclei concentration measured with the Countess-3 Automated Cell Counter. After this, nuclei samples were processed in an identical way as to the cell samples.

One thousand human IEQs from eight different donors (**Additional file 1: Table S1**) were transplanted into the renal sub-capsular space of 4–5-month-old euglycemic RAG1^−/−^ mice as previously described in detail (17,27, 28). Human islet grafts were harvested 1.5-3 months after transplantation, and nuclei were isolated as indicated above. Animal studies and procedures were performed with the approval of and in accordance with guidelines established by the Institutional Animal Care and Use Committee of the Icahn School of Medicine at Mount Sinai (IACUC #2015– 0107).

### Single-cell and single-nucleus RNA sequencing, alignment, and matrix generation

Description of the methods used to obtain cells and nuclei samples for these studies has been already published (17). Briefly, cells and nuclei were prepared according to the 10X Genomics Single Cell 3’ V3.1 Reagent Kit protocol, processed with 10X Genomic Chromium Controller for partitioning and barcoding, followed by the cDNA library generation. Samples were sequenced by NovaSeq 6000 System (Illumina) at the Weill Cornell Medicine, Genomics and Epigenomics Core. FASTQ files were aligned with Cell Ranger V.6.1.1 with Single Cell 3’ V3 chemistry on the 10X Cloud’s pipeline. In the analysis, we included the intronic reads only in the snRNA-seq data with GRCh38-2020-A library. For the human islet graft samples, which were also processed identically as the snRNA-seq dataset from *in vitro* human islets, we included intronic reads as well and used the GRCh38-mm10-2020-A library to distinguish human and residual mouse genes. After the 10X h5 format file was generated, data were analyzed on the R platform with Seurat package V.4.3.0 (29).

### Quality control, integration, and projection

Quality control parameters have been recently described for these studies (17). Briefly, ambient mRNA adjustment was performed using SoupX (20% contamination estimation) (30). Cells with less than a 500 gene count, less than 250 gene varieties, less than 0.8 log10 genes per UMI, and a greater than 20% mitochondrial gene ratio were filtered out. Doublets were algorithmically removed with the Doubltfinder package (31). Human *in vivo* islet datasets were projected onto our integrated scRNA/snRNA-seq dataset as a reference. Since human islet grafts from harvested mouse kidneys naturally contain residual mouse cells and their mRNAs, nuclei with more than 10% of mouse genes were filtered out during quality control process. For the *in vivo* human islet graft data, we used a 10% doublet rate that is appropriate to define several human islet cell populations with cleaner cluster edges and more defined UMAP structures. After data quality control, scRNA-seq and snRNA-seq data were integrated using Seurat’s SCTransform function without allocating method parameters, and cell type identity was assigned according to the normalized gene expression level, referencing the canonical pancreatic cell type genes, as previously published (17,29,32).

### Data processing with α-cell cluster

We previously integrated three donor-matched datasets (single-cell RNA and single-nucleus RNA sequencing) (17) and identified the α cell cluster (*GCG*^high^, *CRYBA2*^high^, *PCSK2*^high^, *TTR*^high^), that was subsequently subset into separate data. We next reassigned cell type identities back to the original numbering generated by the Louvain algorithm with a 0.8 resolution in Seurat’s FindNeighbors() function. This yielded ten distinct clusters; however, five of them were composed of fewer than 10 cells. As such, we grouped these small clusters together with their nearest neighbors.

### Reference based projection

We previously annotated cell types in scRNA/snRNA-seq of human islets *in vitro* (17). In the current study, we updated the cell types with five α- and three β-cell subpopulations. We then set it as a reference, and projected all data investigated in this study including human islet graft snRNA-seq and HPAP-HIRN (see below) adult non-diabetic and T2D human islet scRNA-seq data, in this reference (26).

### Pathway analysis

Single-cell level gene set enrichment analysis was performed to define the molecular and cellular processes using the escape package, which accesses the entire C2 and C5 library from Molecular Signature Database (v.7.0) (33,34). The enrichment score for the entire C2 (6,495 pathways) and C5 (15,937) sets were calculated for each cell. Then we used getSignificance() function from escape package to rank the differentially enriched pathways among subpopulations using the ANOVA test for *fit* parameter due to the multiple groups. Additionally, we selected relevant pathways such as α-cell, glucagon, metabolites, and metabolism with qualifying statistics for analysis and visualization.

### RNA velocity and PAGA analysis

RNA velocity analysis was conducted with integrated sample count tables generated by Matrix package. Velocyto was used to generate a Loom file referencing GRCh38 human genome or GRCh38-mm10 human (35) and mouse genome for relevant data. UMAP coordinates, and α- and β-cell subtype labels were then extracted from the Seurat data to maintain identical UMAP location. An AnnData file was created from the data using scanpy Package (36) and spliced/un-spliced counts were measured by scVelo package with proportions() function (37). A stochastic model hyperparameter was used to estimate velocity, which was visualized with velocity streamline of α- and β-cell sub-clusters transitions. The scVelo algorithm was used to generate PAGA trajectories based on connectivity data (36,37) with the original UMAP coordinates extracted from the Seurat data object.

#### Commonality and splicing analysis

We extracted UMAP coordinate and cell type annotation from the Seurat data frame, then transformed it to the Monocle 3 data object. To calculate the pseudo-temporal gene expression association, we assigned the base according to the PAGA result, then subset the relevant clusters, and calculated the genes with high pseudo-temporal association with graph_test() function. Next, we defined the genes which share the common transitional direction. Subsequently, we obtained spliced/unspliced counts of genes from Anndata data object that contains RNA velocity results and calculated the regression by assigning unspliced count on x-axis, spliced count on y-axis.

### Analysis of non-diabetic and T2D donor islet scRNA-seq data from the Human Pancreas Analysis Program (HPAP) of the Human Islet Research Network (HIRN)

To evaluate α-β cell plasticity changes in T2D, we processed publicly available HPAP-HIRN scRNA-seq data (26) obtained from islets from 13 adult non-diabetic and 13 adult T2D donors using CellRanger (V7.1.0) on the 10X cloud platform, referencing intron-inclusive GRCh38-2020-A transcriptome. To decide whether we should use intron reads or not, we have processed them with and without intron hyperparameter. Then we projected them into our scRNA/snRNA integrated reference, and calculated prediction score of cell type annotation. We observed improved prediction score with data processed with intron reads (0.666 vs. 0.704 for non-diabetic samples, 0.694 vs. 0.753 for T2D samples). Thus, we used data with intron reads for further analysis.

### Statistical analysis

Data are presented as means ± SE in bar graphs, violin plots, scatterplots, and text. Statistical significance analysis was performed using Wilcoxon rank-sum test or Student’s t test for comparison between groups as indicated in the figure legends. P < 0.05 was considered statistically significant. The simplified asterisk statistical significance annotation followed conventional criteria of 0.05, 0.01, 0.0001, and 0.0001 for increment number of asterisks.

### Study approval

All protocols were performed with the approval of and in accordance with guidelines established by the Icahn School of Medicine at Mount Sinai Institutional Animal Care and Use Committee (IACUC #2015–0107).

## RESULTS

### Comparative scRNA-seq and snRNA-seq analysis distinguishes five different α-cell subtypes in human islets *in vitro*

We have previously shown that integrated scRNA-seq and snRNA-seq analysis of human islets can distinguish three different β-cell subtypes with different transcriptome profiles *in vitro* and *in vivo* in human islet grafts (17). The combination of the datasets from both RNA-seq platforms effectively increases the analytical power by providing additional information from both cytoplasmic and nuclear transcriptomes. Using the same dataset and integrated reference, we aimed here to define α-cell subtypes in human islets (**Fig. 1A**). After sub-setting the identified α-cell cluster, we generated sub-clusters with a Louvain resolution of 0.8. We then grouped minor α-cell clusters containing fewer than ten cells into clusters with closest proximity and assigned five α-cell sub-clusters. (**Fig. 1A-C**). Subclusters were named according to gene expression and their UMAP location related to the proximity to a neighboring β-cell cluster (β1, β2, β3, from 17): α1, α2, α-β-transition 1 (AB-Tr1, proximity to β1), α-β-transition 2 (AB-Tr2, proximity to β2), and α-β (AB) cluster **(Fig. 1D**). We employed this unbiased strategy instead of marker-based subcluster assignment since it considers both most of the transcriptome and the multiple PCA dimensions assigned to calculate clusters. The total number of cells analyzed per subcluster and transcriptomics platform in the three human islet preparations appear in **Suppl. Fig. 1A.** The α1, and α2 sub-clusters comprise most α-cells (68-73%) and are similar between scRNA- and snRNA-seq datasets. In contrast, AB-Tr2 and AB clusters are different in proportion between scRNA- and snRNA-seq datasets (**Fig. 1E**), with a larger proportion of AB-Tr2 cells in scRNA-seq than in snRNA-seq (20% vs. 8.6%), and fewer AB cells (2.4% vs. 6.4%). This suggests that snRNA-seq with pre-mRNA analysis can uncover more multihormonal cells (see below) than scRNA-seq.

**Figure 1.**
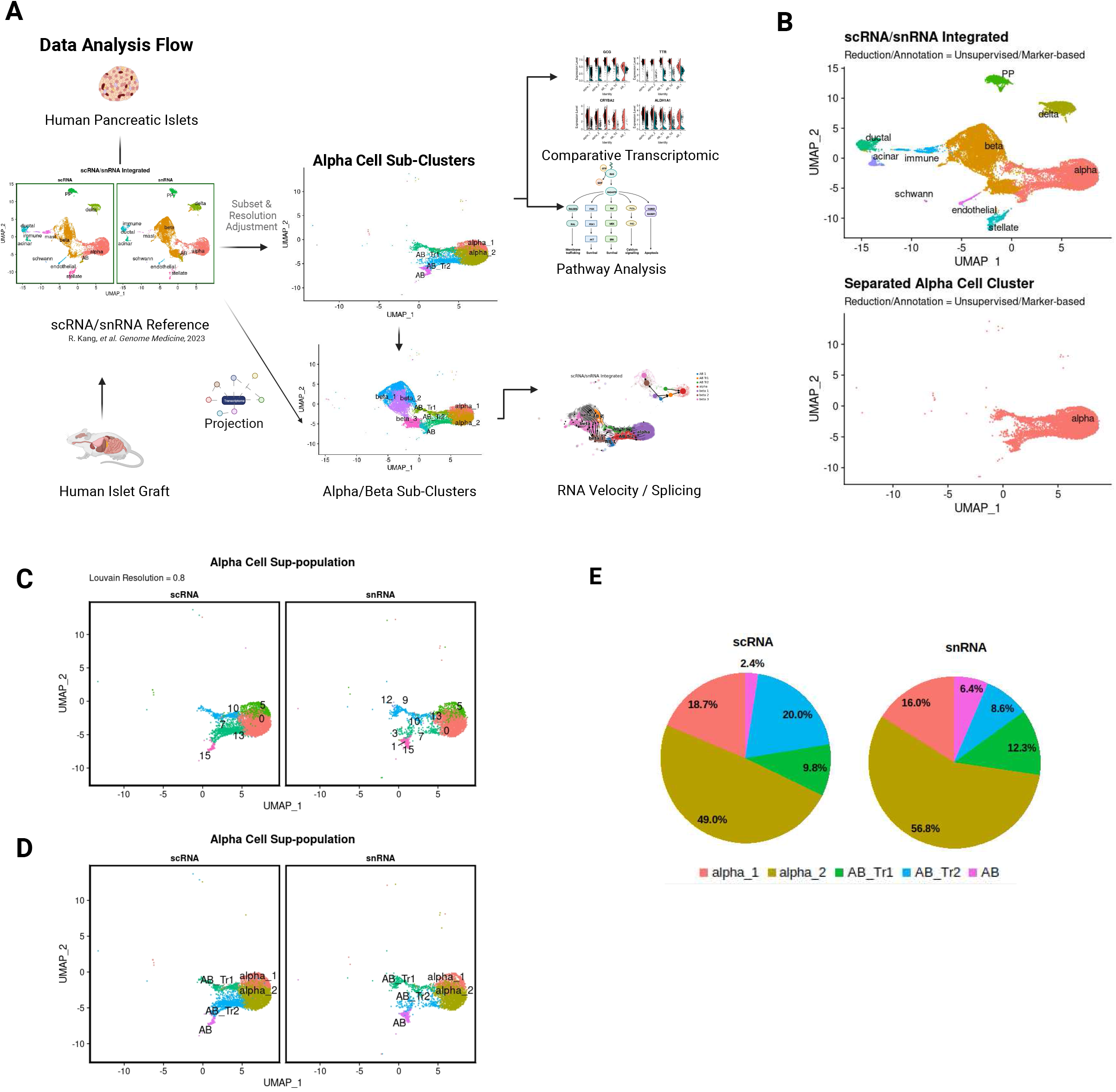
Experimental Design, Unsupervised Clustering, and Sub-Clustering of Alpha Cells. **A.** Human islet processing and data generation scheme. **B.** Unsupervised clustering, cell type annotated UMAP and separated α-cell cluster (below). **C.** Pre-annotated α-cell subclusters by assigning Louvain resolution 0.8; and **D.** Annotated α-cell clusters according to the UMAP location relative to neighboring β-cells and gene expression. UMAP is split by the processing type – scRNA (left) and snRNA (right). **E.** Proportion of the α-cell subtypes in scRNA and snRNA-seq datasets.

### Signature gene expression pattern of the human islet α-cell subtypes *in vitro*

**Fig. 2A** compares expression of canonical alpha cell genes in human islets in scRNA-seq and snRNAseq datasets. Expression of *GCG* and *ALDH1A1* genes was high across the five α-cell subclusters and was similar in scRNA- and snRNA-seq datasets. In contrast, *TTR* and *CRYBA2* showed substantially higher expression levels in scRNA-seq than in snRNA-seq datasets, where their expression was marginally detectable or absent in some α-cell subtypes (**Fig. 2A**). This provides further support for our previous observation that the *GCG, CRAYBA2, ALDH1A1, TTR* gene set optimally defines α-cells in scRNA-seq analysis of human islets but is suboptimal for snRNA-seq annotation (17). Therefore, we next analyzed the expression of our recent α-cell gene set derived from snRNA-seq (17) and observed that *PTPRT, FAP, PDK4* and *LOX4* show higher expression in clusters in the snRNA-seq than in the scRNA-seq dataset (**Fig. 2B**). Interestingly, these four latter genes showed a decreasing pattern from α1 to AB cells.

**Figure 2.**
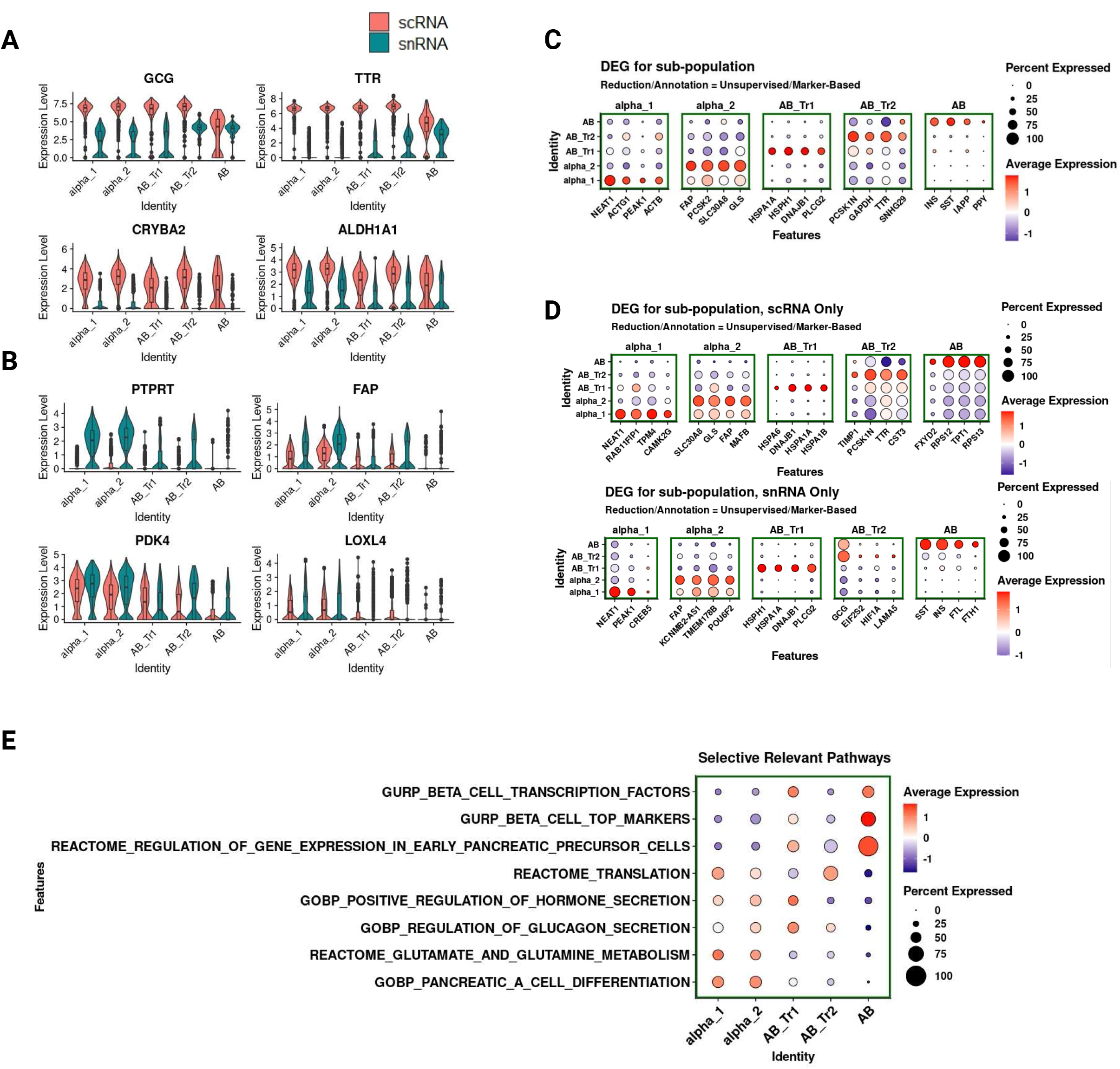
Gene expression, DEG, and Pathway Enrichment Analysis. **A.** Gene expression of 4 canonical α-cell markers, split by processing types. **B.** Gene expression of single nucleus markers previously identified (17). **C.** Dot-plot visualization of top 4 differentially expressed genes. **D.** Differentially expressed genes for α-cell sub population in scRNA data (top) and snRNA data (bottom). **E.** Pathway enrichment analysis for α-cell subpopulations. We searched pathways using keywords for pancreatic endocrine cells (α/β/δ/PP/ε), glucagon and hormone signaling/processing/secretion, and cellular development (differentiation, precursor, dedifferentiation, development, senescence) and arrange the pathway order according to the enrichment patterns from α1 to AB cells.

To better define gene sets that identify the different α-cell subclusters and their transcriptome differences, we performed differential gene expression (DEG) analysis for every cluster against all the remaining clusters. As expected, canonical genes (*GCG, TTR, LOXL4* and *CRYBA2*) or the previously defined snRNA-seq gene markers (*PTPRT, PDK4*) do not appear among the top differentially expressed genes in the α-cell subclusters despite of the qualifying p-value and fold changes (not shown). However, we found considerable differences in the transcriptome among the five different α-cell subclusters (**Fig. 2C**). Using the integrated scRNA- and snRNA-seq dataset, we found that the α1 subcluster displayed selectively higher expression of *NEAT1, ACTG1, PEAK1* and *ACTB*. The α2 subcluster favored *FAP, PCSK2, SLC30A8,* and *GLS expression.* The AB-Tr1 subcluster expressed *HSPA1A, HSPH1, DNAJB1,* and *PLCG2* most highly. AB-Tr2 subcluster favored *PCSK1N, GAPDH, TTR,* and *SNHG29*. And the AB cell subcluster displayed expression of pan-endocrine hormonal genes such as *INS, SST, PPY* and the β-cell identity gene *IAPP* (**Fig. 2C, Suppl. Fig. 1B**). Interestingly, most of AB differentially expressed genes were defined by the snRNA-seq which employs intron-inclusive references. This suggests that a large portion of these genes were pre-mRNA, as might be expected of RNA collected from the nuclear compartment. (**Fig. 2D**). Due to the similar transcriptome among subclusters, some of the genes appeared in two clusters. We therefore substituted the next best DEG to avoid duplicated charts for different subclusters. Importantly, while this DEG analysis is useful to understand the differences among subclusters, their functional significance in a specific subcluster is uncertain. Therefore, we performed pathway analysis of the five different human islet α-cell subclusters.

### Pathway analysis of α-cell subclusters in human islets *in vitro* reveals potential different functions

In an attempt to bridge the transcriptome of the five alpha cell subtypes to possible α-cell subcluster-specific functions, we performed single-cell-level pathway enrichment analysis using the integrated scRNA- and snRNA-seq dataset (17), the Msigdb database (33), and the escape package (34). We targeted gene set enrichment analysis of common pathways in pancreatic α-cells and screened for glucagon production and secretion, metabolism, differentiation, and translation-related pathways, then selected the pathways with qualifying p-value and FDR. As expected, we observed differential enrichment of pathways among the various α-cell subclusters (**Fig. 2E**). While α1 and α2 cell subclusters displayed higher enrichment for α-cell differentiation and glutamate/glutamine metabolism pathways, AB-Tr1 cells showed enrichment for glucagon secretion (also present in α1 and α2 cells albeit at a lower level). α1 and AB-Tr2 demonstrated the highest enrichment for protein translation-related pathways, while the AB cluster displayed the highest enrichment for early pancreatic precursor cell gene expression, top β-cell markers, and β-cell transcription factors (**Fig. 2E**).

Non-targeted DEG analysis of each α-cell subcluster revealed the top 4 pathways for each subcluster according to the p-value and FDR (**Suppl. Fig. 1C**). The α1 subcluster was enriched by pathways exemplified by vesicle transport, cell adhesion, TCA cycle, and carbohydrate catabolic processes. Presumably, the function of this cluster is related to carbohydrate metabolism and cell-cell communication. The α2 subcluster was enriched for neuron and neurotransmitter-related pathways, seemingly suggesting a neuroendocrine-responsive cluster. Interestingly, it also showed high enrichment for the mitotic cell cycle process (**Suppl. Fig. 1C**). The AB-Tr1 cell subcluster displayed enrichment in NAD metabolic process, the inclusion body assembly, and acetylcholine response related pathways. Additionally, it was the only cluster that was enriched in genes that define the trans-differentiation pathway (exemplified by *PDX1, INSM1* and *SMAD3*), although the enriched cell population was small (∼25% cells) (**Suppl. Fig. 1C**). The AB-Tr2 cell subcluster was highly enriched with ribosome-related pathways suggesting enhanced protein translation capability. Interestingly, it was also enriched with polyamine transport (**Suppl. Fig. 1C**). Finally, the AB cell subcluster, despite its clear α-cell phenotype, also displays high *INS*, *SST* and *PPY* gene expression. This multihormonal cell subcluster with a topographical location very near to the β-cells, exhibited high enrichment for neurotransmitter/noradrenergic neuron differentiation, calcium ion regulation-related pathways such as postsynaptic cytosolic calcium ion concentration and intracellular calcium activated chloride channel activity (**Suppl. Fig. 1C**).

### RNA velocity and PAGA analysis of α-β-cell trajectory inference in human islets *in vitro* using the integrated scRNA- and snRNA-seq reference

We next analyzed RNA velocity using Scanpy and scVelo packages to investigate a possible transition among α- and β-cell subclusters. The α-cell subclusters identified in this study were combined with the β-cell subclusters that we have previously reported (17). From the integrated reference dataset, we observed a comparable ratio of spliced/unspliced RNA among the different cell subtypes, except for the AB-Tr2 subcluster, which had the most dominant proportion of spliced mRNA (**Fig. 3A**) and the highest enrichment in ribosome-related pathways and translation (**Suppl. Fig 1C**), perhaps suggesting that the AB-Tr2 subcluster is enriched for mRNA splicing and protein translation. The β1 subcluster (most mature β-cell subcluster, 17) had the most dominant proportion of unspliced mRNA (**Fig. 3A**). We then visualized the streamlined velocity plot. Here, we observed general bifurcating pattern from the AB/β3 cluster both towards the β1 and α2 subcluster (**Fig 3B**). Using velocity length to characterize speed of transition or differentiation, both the β2 and the AB-Tr2 clusters showed significantly higher length than the other clusters, denoting vibrant splicing activity and satisfying overall velocity confidence across all the α- and β-cell subpopulations (**Figure 3C**). Additionally, we measured the velocity of several representative genes of α- and β-cells and found that the expression level and velocity did not always correlate with each other. *GCG* expression was highest in the transitional clusters (AB_Tr1/AB_Tr2), but RNA velocity was higher in α2 and part of the α1 subcluster. *INS* expression level was selectively higher in β-cells, while RNA velocity displayed slight induction in the AB and AB_Tr2 subclusters. We also assessed the β-cell gene, *GLP1R,* and found that its expression pattern matched with RNA velocity. Similarly, we also assessed the α1 cell specific gene, *NEAT1,* and found that it showed higher expression in β1 but very low RNA velocity (**Fig. 3D**). Together, these observations support the notion that cells in the β3 and AB subclusters may spontaneously transition to more mature β- and α-cell types.

**Figure 3.**
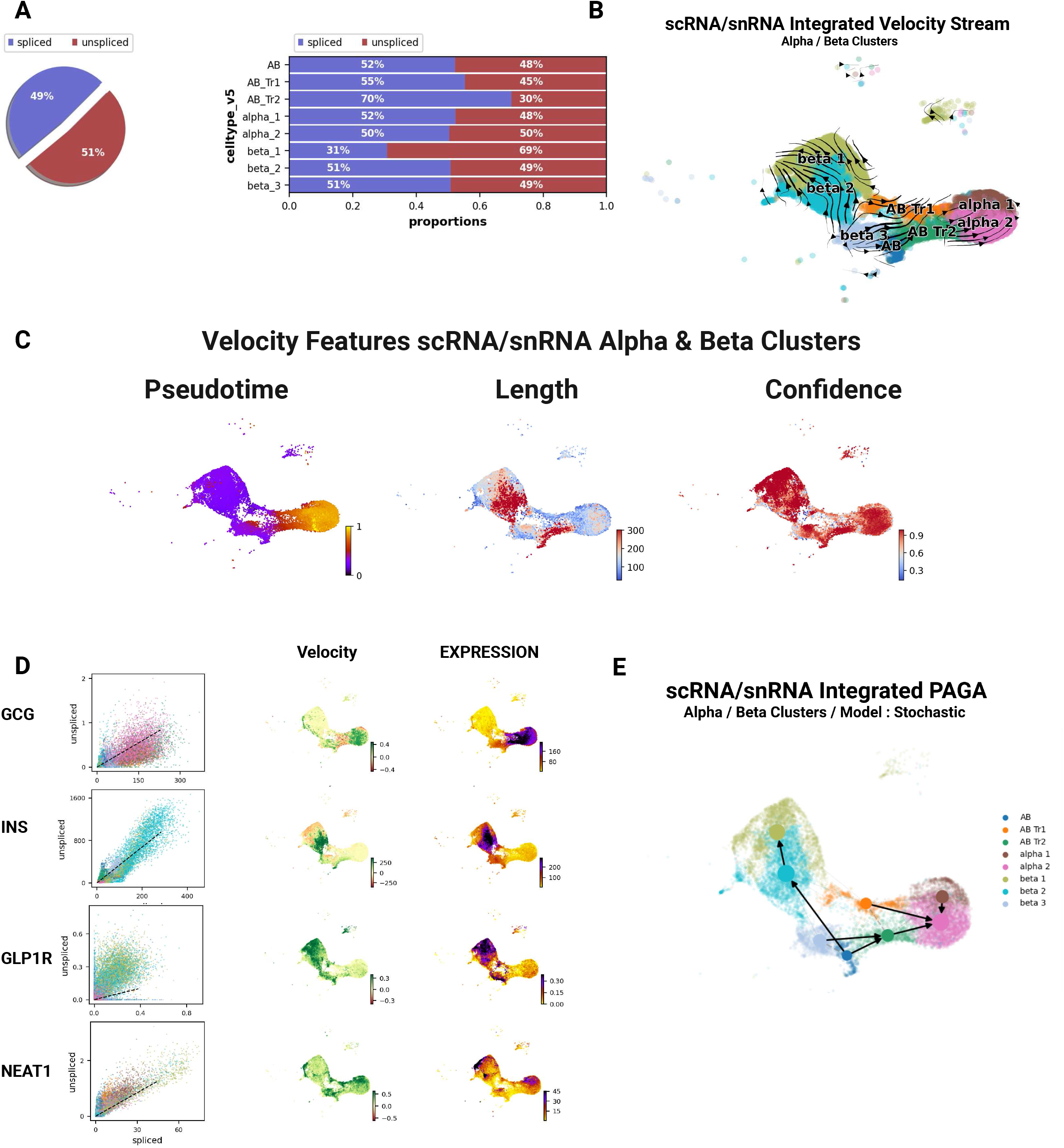
Velocity and Splicing in scRNA and snRNA *In Vitro* Data and Trajectory. **A.** Splicing ratio in α-and β-cell subclusters. **B.** Velocity streamline with scVelo. Stochastic modeling was used. **C.** Velocity length (top) and velocity confidence (bottom). **D.** Single gene velocity visualization with spliced/unspliced scatter plot (left), RNA velocity (mid), gene expression (right) for *GCG, INS, GLP1R* and *NEAT1* **E.** PAGA trajectory analysis based on stochastic modeled velocity.

We next performed PAGA trajectory analysis on this dataset, from which we inferred four main trajectories among α- and β-cell subpopulations: 1) AB-β3-β2-β1 (Trajectory A) which shows AB cells progressing towards the high insulin-secreting β1 subtype; 2) β3-ABTr2-α2 (Trajectory B), showing transition from the less mature β-cell subtype (17) to mature α-cells; 3) AB-ABTr2-α2 (Trajectory C), from multihormonal cell cluster to mature α-cells; and, 4) ABTr1-α2 (Trajectory D) from less differentiated α-cell cluster to a more differentiated α2-cell cluster (**Fig. 3E and 4A**). Subsequently, we investigated the high pseudo-temporal associated gene sets for each trajectory (**Fig. 4B-E**). First, we transformed the data set into the Monocle3 format and then assigned the root as the unbiased PAGA origin for pseudo-time scoring. We ranked the gene list according to Moran’s I score for visualization and clustered the genes according to the patterns (**Fig. 4B-E**). For the transition from AB to β1 (Trajectory A), there were 2 main transitional patterns: increase or biphasic (increase then decrease) (**Fig. 4B**). The transcriptome in the increased gene pattern from AB to β1 is involved in cellular ion regulation and signaling (*TRPM3, DPP6, CASR, KCNMA1*) and insulin release like *ABCC8*. Moreover, neural development and synaptic function genes (*LSAMP, DLG, NRG1*) are also increased in this trajectory. The genes with a biphasic pattern of expression are related to ribosomal protein genes (RPLs, RPSs) and ferritin proteins *FTL* and *FTH1* that are related to iron movement. (**Fig. 4B**). In contrast, the transition from β3 to α2 (Trajectory B) showed increasing expression pattern of apoptosis and proliferation-involved genes (*KIAA1324, BTG2, MALAT1, NR4A2*), glucagon-related genes (*PCSK2, ARFGEF3*) and glucose homeostasis-related genes (*SLC30A8, PAPPA2*) (**Fig. 4C**). Unlike Trajectory A, Trajectory B showed increasing patterns of expression of ribosomal protein genes (RPLs, RPSs) and metabolism-associated transcriptomes (*SLC7A2, PDE10A, GLS*) and decreased expression of β-cell identity genes (*IAPP, INS*) (**Fig. 4C**). In the transition from AB to α2 (Trajectory C), β-cell identity genes (*IAPP, INS*) showed decreasing pattern of expression (**Fig. 4D**) while apoptosis and proliferation-involved genes (*TM4SF4, MALAT1, NR4A1, NR4A2, BTG2, KIAA1324*), ribosomal protein genes, and metabolism and cellular signaling genes (*GLS, SLC30A8, PDE10A*) were increased (**Fig. 4D**). Finally, AB-Tr1-α2 transition (Trajectory D) displayed increased expression pattern of cellular transport and metabolism (*SLC7A2, ALDH1A1*), glucose regulation (*ARFGEF3, SLC30A8, PAPPA2, PTPRN2*), and development/differentiation *(PLCE1, NR4A1, KIAA1324, TM4SF4*) genes. On the other hand, β-cell identity genes (*INS, IAPP, ZNF385D, TRPM3*) displayed reduced expression (**Fig. 4E**).

**Figure 4.**
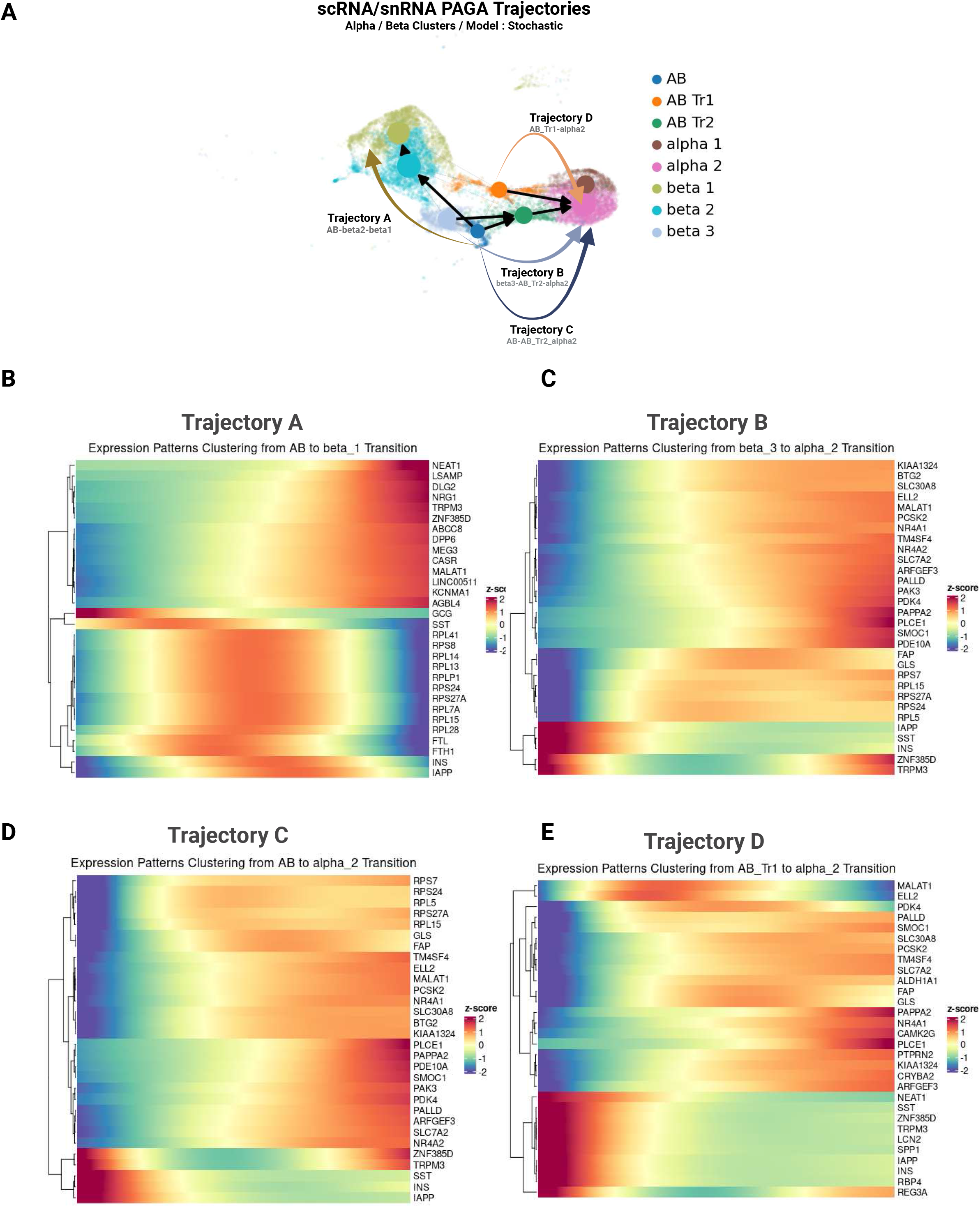
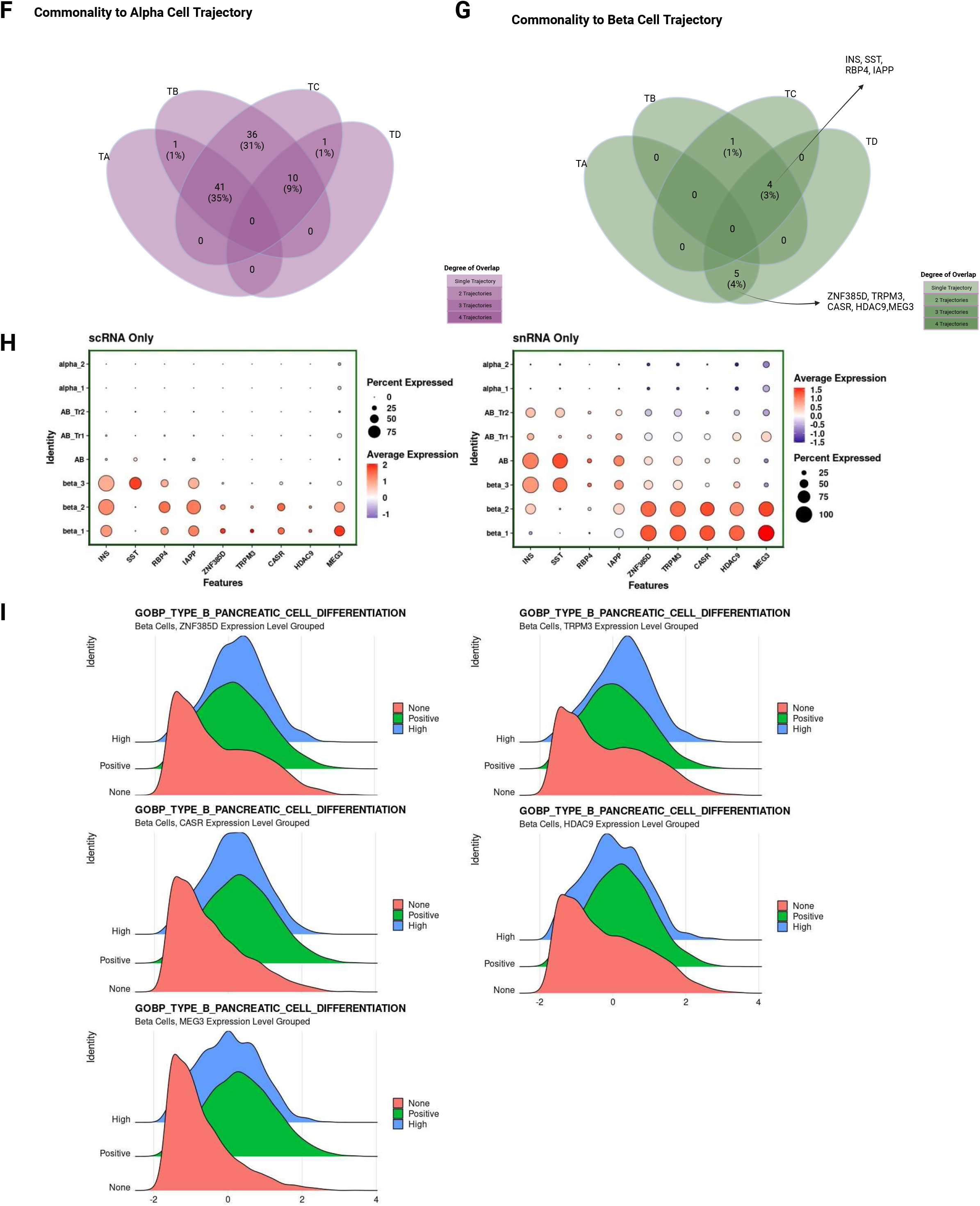
Trajectory and Associated Gene Expression. **A.** Diagram of 4 main trajectories from PAGA analysis; Trajectory A – from AB to β1 (toward β-cells); Trajectory B- from β3 to α2 (toward α-cells); Trajectory C – from AB to α2 (toward α-cells); Trajectory D – from AB_Tr1 to α2 (toward α-cells). **B.** Expression of genes with selective high association with Trajectory A (from AB to β1). Genes were clustered according to the expression pattern. The visualization is done by Monocle3, by separating the relevant clusters and assigning the root according to the PAGA analysis. **C.** Expression of genes with selective high association with Trajectory B (β3 to α2). **D.** Expression of genes with selective high association with Trajectory C (AB to α2). **E.** Expression of genes with selective high association with Trajectory D (AB_Tr1 to α2). **F.** Gene commonality analysis towards α-cell transition. Genes in decreasing pattern in Trajectory A and increasing pattern in Trajectories B, C, and D were intersected leading to common genes among different trajectories. **G.** Gene commonality analysis towards β-cell transition. Genes in increasing pattern in Trajectory A and decreasing pattern in Trajectories B, C, and D were intersected leading to common genes among different trajectories. **H.** Dot plots representing gene expression of the commonality genes between reverse-Trajectory C and D (decreasing pattern of expression) - *INS, SST, RBP4* and *IAPP* and Trajectory A and reverse Trajectory D (decreasing pattern) - *ZNF385D, TRPM3, CASR, MEG3* and *HDAC9* in scRNA-seq (left) and snRNA-seq (right). **I.** Enrichment ridge plot of GOBP’s pancreatic β-cell differentiation pathway. We grouped β-cells in three groups by single gene expression level for each gene (*ZNF385D, TRPM3, CASR, MEG3* and *HDAC9*) and grouped them into three groups – negative (0 normalized expression), positive (0 to 98%) and high (98% and above).

### Gene commonality analysis in trajectory inference

To identify common genes in the four trajectories that can define the transition among different cell types, we performed gene commonality analysis (**Fig. 4F-G**). We divided gene expression patterns into two groups: AB-to-β cells (Trajectory A) and β3/AB/AB-Tr1-to-α cells (Trajectories B, C, D). Common α-cell trajectory genes from AB/β/AB-Tr1-to-α cells (trajectories B, C, D) were S*MOC1, PLCE1, PAPPA2, ZNF331, ALDH1A1, SLC30A8, BTG2, TM4SF4, NR4A1* and *PSCK2* all related to cell signaling, metabolism, development, and growth. On the other hand, β cell trajectory genes from AB-to-β cells (Trajectory A) were *ZNF385D, TRPM3, CASR, MEG3* and *HDAC9*, shared by Trajectory A and inverse Trajectory D; and *INS, SST, RBP4, IAPP* shared by inversed trajectories C and D. **(Fig. 4G).** All these genes, except *SST*, are upregulated in mature more functional β1 and β2 cells while their levels are decreased in β3, AB and the rest of α-cells (**Fig. 4H**), suggesting that alteration in the levels of *ZNF385D, TRPM3, CASR, MEG3* and *HDAC9* can induce changes in the transition from AB cells to β-cells. Interestingly, the highest expression of *ZNF385D, TRPM3, CASR, MEG3* and *HDAC9* occurs mostly in the snRNAseq where pre-mRNA expression and processing might be of great importance to determine cellular trajectories towards transition to different states. Importantly, these data also illustrate the concept that only by integrating scRNA- and snRNA-seq datasets can one visualize and identify key genes in the cellular trajectories from multihormonal AB cells to β-cells. Finally, the degree of expression of these genes positively correlates with GOBP pathway of pancreatic β-cell differentiation (**Fig. 4I**).

### snRNA-seq analysis of α-cell subtypes in human islet grafts

We next turned our attention from *in vitro* gene sets discussed up to this point, to comparable data from our earlier publication (17), obtained from healthy human islets, living, and functioning *in vivo*. Specifically, we determined whether the α-cell subtypes identified in the scRNA-seq/snRNA-seq analysis of isolated human islets from non-diabetic donors (*in vitro* study) were present in human islets in islet grafts (*in vivo* study). We projected the snRNA-seq dataset from human islet grafts transplanted in immunosuppressed mice onto the *in vitro* scRNA-/snRNA-seq dataset reference which is pre-labeled with α-cell subcluster information (**Fig. 5A**). To our surprise, the projection algorithm failed to recognize the AB subcluster in the human islet graft *in vivo* snRNA-seq dataset: only α1, α2, AB-Tr1 and AB-Tr2 were present (**Fig. 5B**). Together, α1 and α2 cells comprised close to 96% of the α-cells observed *in vivo* in contrast with the 68-73% of these cells found *in vitro* (**Fig. 5C** vs. **1E**). Comparison of α-cell subtype proportions between *in vitro* and *in vivo* shows a remarkable significant increase in α2 cells up to 92.5% while the abundance of other α-cell subtypes significantly decreased (**Fig. 5D**). This indicates that the *in vivo* environment favors α2 cells (the most differentiated α-cell subtype), while AB cells (the least differentiated α-cell with bifurcated α/β-cell transitional properties), disappear in the human islet grafts. This raises the possibility that AB cells may reflect a non-physiologic response to islet isolation and culture.

**Figure 5.**
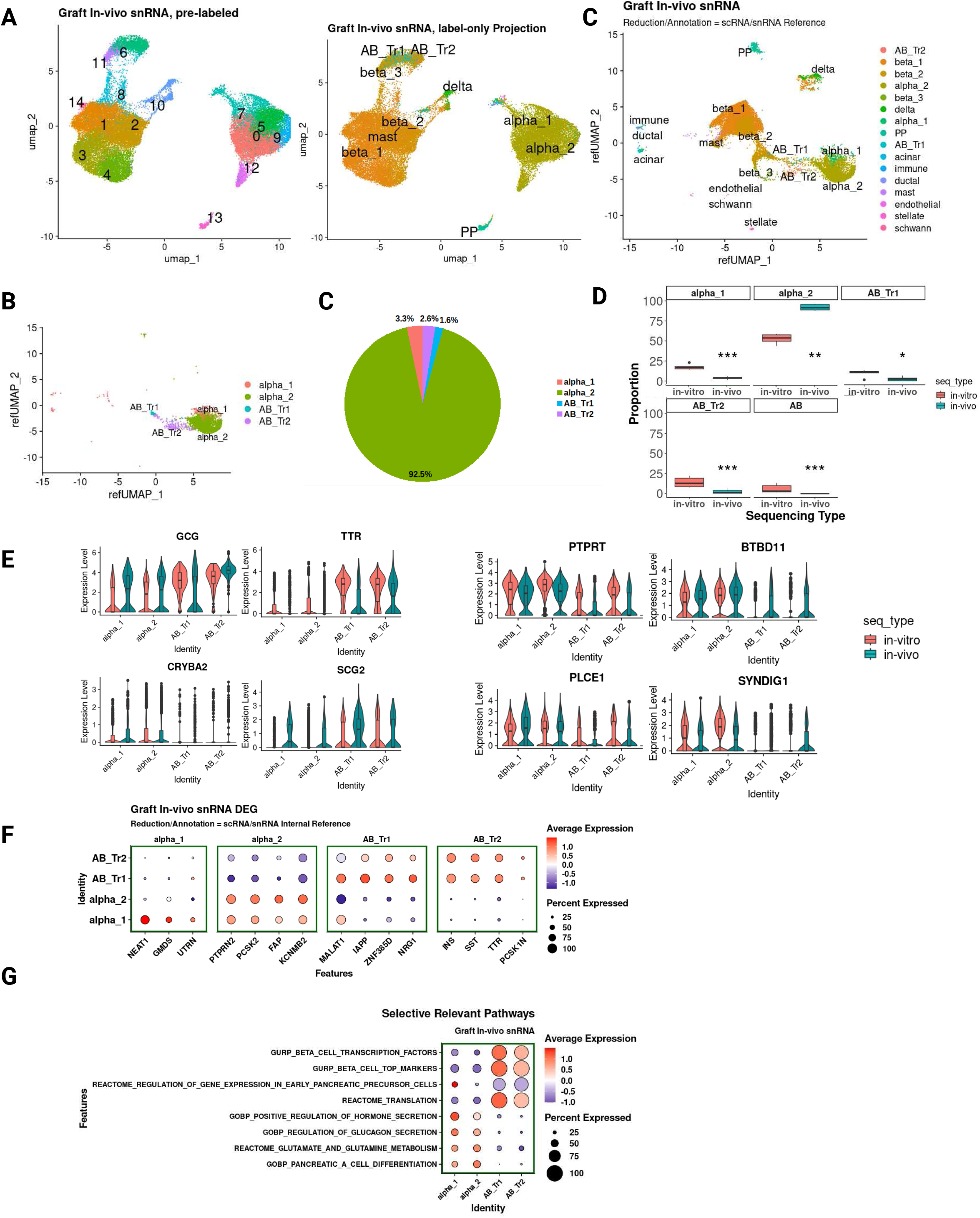
Clustering, Projection and Gene Expression in Human Islet Grafts *In Vivo.* **A.** Unsupervised clustering of human islet grafts *in vivo* snRNA data. Numbers are automatically generated by Seurat neighboring algorithm (left). Label-only transferred UMAP (mid) in which the cell type annotation is from *in vitro* data. Human islet graft *in vivo* snRNA was projected on *in vitro* scRNA/snRNA reference both for cell type annotation with α-and β-subclusters and UMAP embedding (right). **B.** Separated α-cell cluster retaining α-cell subcluster annotation and UMAP coordinate from *in vitro* reference. **C.** α-cell subtypes proportion, **D.** α-cell subtypes proportion comparison between *in vitro* and *in vivo* data and analyzed statistically by Wilcoxon rank-sum test. The proportion of cells of each donor was used to calculate statistical mean differences. **E.** Comparative transcriptomic data between *in vitro* and *in vivo* for four canonical α-cell markers (left), and the four newly defined single nucleus α-cell markers (17) (right). **F.** Differentially expressed genes in each subcluster in the *in vivo* snRNA data **G.** Pathway enrichment analysis for *in vivo* α-cell subpopulations. We applied an identical strategy to identify the relevant pathways as Figure 1E.

Expression of canonical and newly identified genes in the α-cell subclusters in the snRNA-seq data of human islet grafts follow a similar pattern of expression compared with snRNA-seq in human islets *in vitro* (**Fig. 5E**). Interestingly, we observed that hormone genes (*SST, INS*, and *PPY*) which were selectively higher in the AB cluster *in vitro*, are now present in *GCG*-expressing AB-Tr1 and AB-Tr2 (**Fig. 5F**). We speculate that the *in vivo* microenvironment might induce transcriptome changes that lead to the migration of AB cells into transitional clusters, resulting in the localization of *GCG-, INS-* and *SST-*positive cells in the AB-Tr1 and AB-Tr2 clusters. On the other hand, α2 cells selectively expressed the 4 top genes identified in human islets *in vitro PTPRN2, HS6ST3, SIPA1L3,* and *KCNMB2* and the AB-Tr2 cells have differentially expressed genes that were also expressed in AB-Tr1 cells, confirming their proximity in transcriptome (**Fig. 5F**).

To understand potential functional differences among α-cell subclusters in human islet grafts, we performed gene set enrichment analysis (**Fig. 5G**). α1 and α2 cells, most α-cells *in vivo*, aligned with pathways mainly related to glucagon secretion and α-cell differentiation similarly to those observed *in vitro*. However, *in vivo* AB-Tr1 and AB-Tr2 subclusters show higher enrichment in pathways related to translation, pancreatic precursor cells and β-cells, as compared to their counterparts *in vitro* (**Fig. 5G**). Additional pathways for each cell type are shown in **Suppl. Fig. 2**.

### RNA Velocity and PAGA analysis of α-β-cell trajectory inference in human islet grafts using the integrated scRNA- and snRNA-seq reference

As expected of RNAs derived from the nuclear compartment, most of the mRNAs in the snRNA-seq data in the human islet grafts are unspliced, although the transitional clusters (AB-Tr1 and AB-Tr2) and the less differentiated β-cells (β3) showed a modestly higher spliced ratio (**Fig. 6A**). Intriguingly, the velocity pseudotime, length and confidence displayed markedly different patterns in the *in vivo* human islet grafts as compared to human islets *in vitro* (**Figs. 6B** vs**. 3C**). In the human islet graft, β3 cells and a small portion of α2 cluster demonstrated longer velocity length (**Fig. 6B**) whereas β2 and AB-Tr2 showed the longer velocity *in vitro* (**Fig. 3C**). This may reflect the different environments *in vivo* vs. *in vitro,* or the fact that *in vivo* data only contains snRNA-seq while the *in vitro* data is an integration of scRNA- and snRNA-seq data.

**Figure 6.**
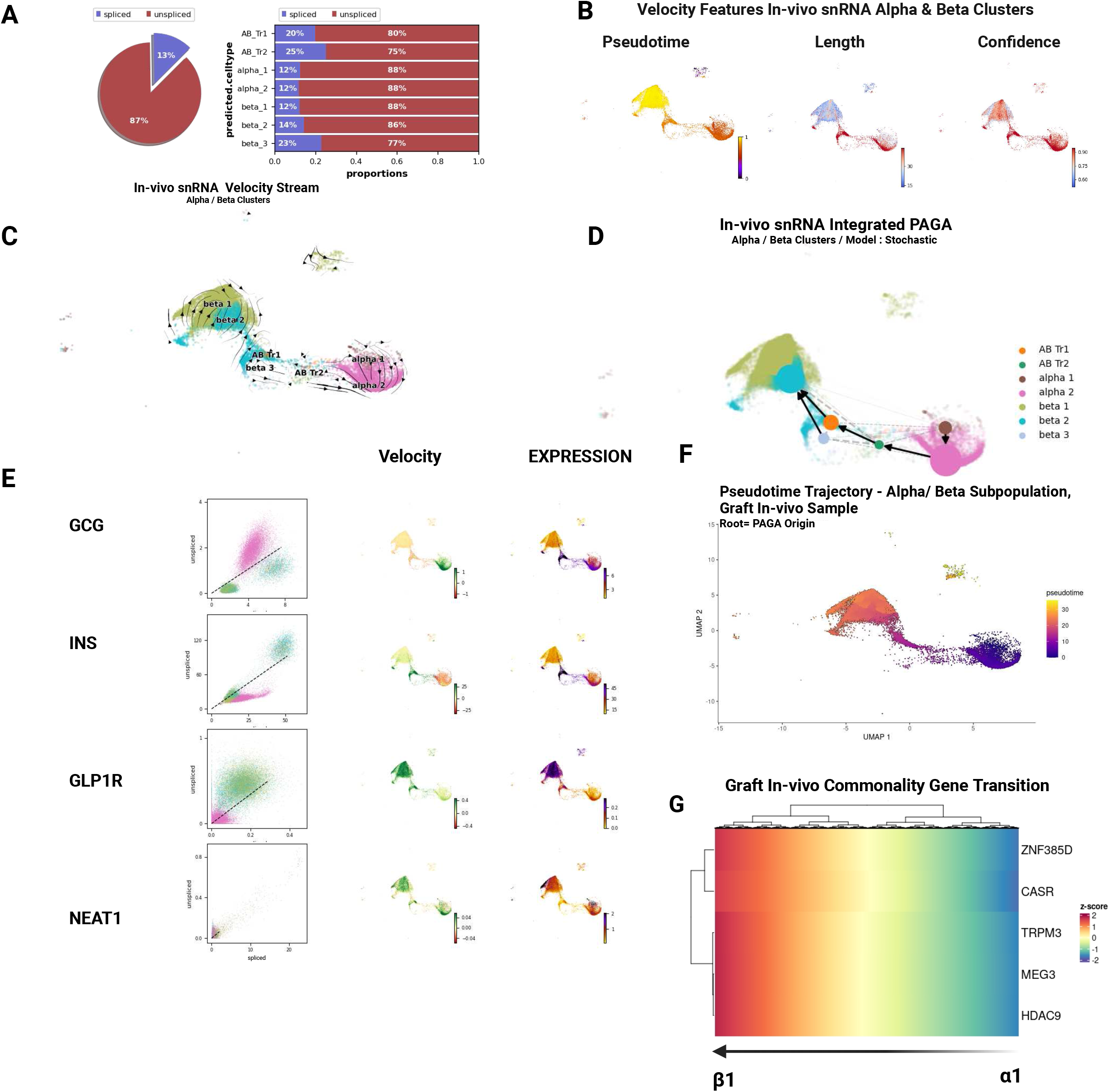
RNA Velocity and PAGA Trajectory Analysis of Human Islet Grafts *In Vivo* snRNA- seq Dataset. **A.** Splicing ratio for α-and β-cell subpopulations. **B.** Velocity-based pseudotime, length and velocity confidence. The analysis was done with stochastic modeling. **C.** Velocity stream from scVelo analysis. The UMAP coordinate and cell type annotation was done by projecting data to scRNA/snRNA *in vitro* reference. **D.** PAGA trajectory analysis. **E.** Single gene velocity visualization with spliced/unspliced scatter plot (left), RNA velocity (mid), gene expression (right) for *GCG, INS, GLP1R* and *NEAT1*. **F.** Pseudotime UMAP of human islet grafts *in vivo* snRNA-seq data. The root was chosen according to the PAGA analysis. **G.** Gene expression of five commonality genes along the pseudotime trajectory (*CASR, ZNF385D, TRPM3, MEG3* and *HDAC9*)

The *in vivo* RNA velocity stream plot and PAGA analysis also differed markedly from their *in vitro* counterparts. Here, we observed a general unidirectional transition from differentiated α-cells α1 and α2 to AB-Tr1, ABTr2 and then to β-cell types but no evidence of transition from β-cells to α- cells as we observed in human islets *in vitro* (**Fig. 6C-D** vs. **3E**). This may suggest that under normal physiologic circumstances, transition of cells from one phenotype to another only occurs from β-cells to α-cells. In contrast, the opposite trajectory may appear only after isolation and culture of human islets. This is reminiscent of the apparent plasticity of α-cells into β-cells when β-cells are obliterated, but not the opposite direction when α-cells are obliterated (18,38).

We next conducted a comprehensive analysis of single-gene velocity in the *in vivo* study for the four specific genes also studied *in vitro*: *GCG, INS, GLP1R,* and *NEAT1* (**Fig. 6E and 4D**). Interestingly, while the *in vitro* dataset exhibited notably elevated *INS* velocity and expression within the β2 cluster, the human islet graft data displayed significantly higher velocity and expression within the β3 and transitional clusters AB-Tr1 and AB-Tr2 (**Fig. 6E and 4D**). On the other hand, *GCG* and *GLP1R* genes closely resembled the patterns observed in the *in vitro* dataset (**Fig. 6E and 4D**). Specifically, *GCG* exhibited heightened velocity and expression within the α2 cluster, while *GLP1R* displayed robust expression within the β subclusters. *NEAT1* demonstrated rapid velocity within the α2 subcluster but exhibited higher expression levels within the α1 subcluster, which contrasted with the *in vitro* dataset where higher velocity and expression were observed in the α1 subcluster (**Fig. 6E and 4D**). These studies revealed distinct patterns of RNA velocity and expression, particularly revealing significant disparities in the behavior of *INS* and *NEAT1 in vivo* as compared to *in vitro*.

We next performed gene expression analysis along the pseudo-temporal transition using Monocle3. We assigned the α1 cluster as the root, arranged the cells by pseudo-time score and ranked the genes with spatial association Moran’s I score as we had for scRNA/snRNA *in vitro* data (**Fig. 6F-G**). We observed a decrease in the expression of several neural development/metabolism genes (*GLS, SYNDIG1, TENM2, POU6F2*) and growth/development related genes (*FAP, PAPPA2, GPC6*) in the α-to-β-cell transition. On the other hand, we observed induction of β-cell metabolism/homeostasis regulator genes (*PLUT, CASR, GLIS3, LDLRAD4, KIRREL3, TRPM3*), cell signaling/neural synapse-related genes (*LRFN2, UNC5D, DOCK10, NRG1*), and genes involved in gene expression regulation and transcription factors (*ZNF385D, HDAC9, MEG3, DACH2, SYNE2, MEG8*) (**Fig. 6F-G** and **Suppl. Fig. 3**).

### α-β-cell trajectory inference in human islets from T2D donors

To evaluate α-β-cell plasticity in T2D human islets, we interrogated the publicly available HIRN-HPAP database and processed scRNA-seq data from islets isolated from non-diabetic and T2D adult human donors (26). We repeated identical data analysis strategies – projection to a scRNA-/snRNA-seq reference, extracting UMAP coordinates, cell type annotation and RNA velocity analysis. Analysis of total RNA splicing ratios within distinct α-β-cell clusters showed that the AB subcluster exhibited a substantial reduction in spliced mRNA, while the AB-Tr1 subcluster displayed the opposite trend (**Fig. 7A**). Furthermore, the α1, α2, and β2 subclusters also demonstrated a notable decrease in overall RNA splicing activity, while the remaining subclusters exhibited either minimal changes or remained largely unchanged (**Fig. 7A**). To compare non-diabetic and T2D datasets, we established a scale ranging from 0 to 600 for velocity length measurements. This allowed us to discern significant differences between the two datasets. Specifically, our findings revealed a distinct pattern in the T2D dataset, with a selective increase in velocity length observed within the AB-Tr1, ABTr2, AB and β3 subclusters (**Fig. 7B**).

**Figure 7.**
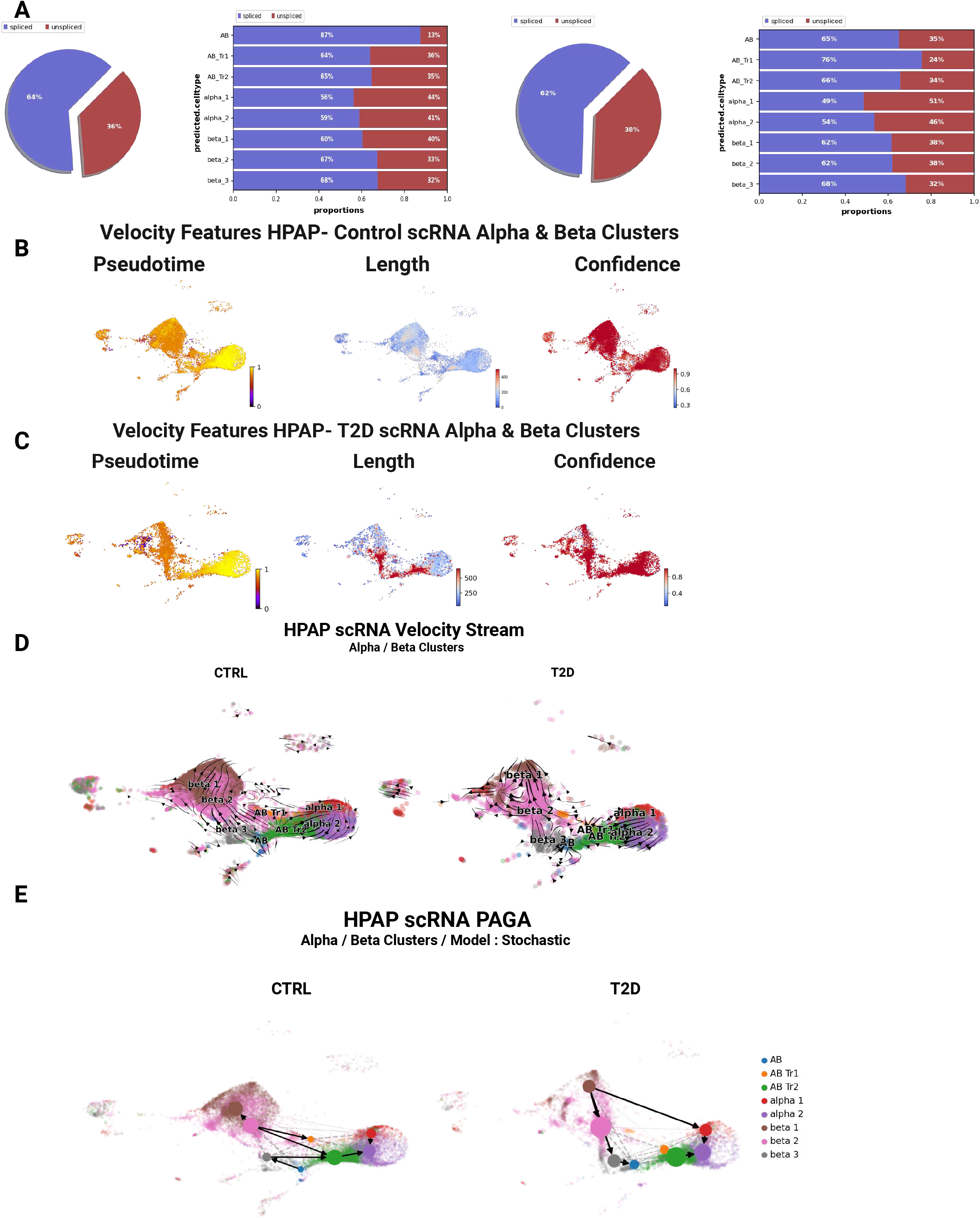
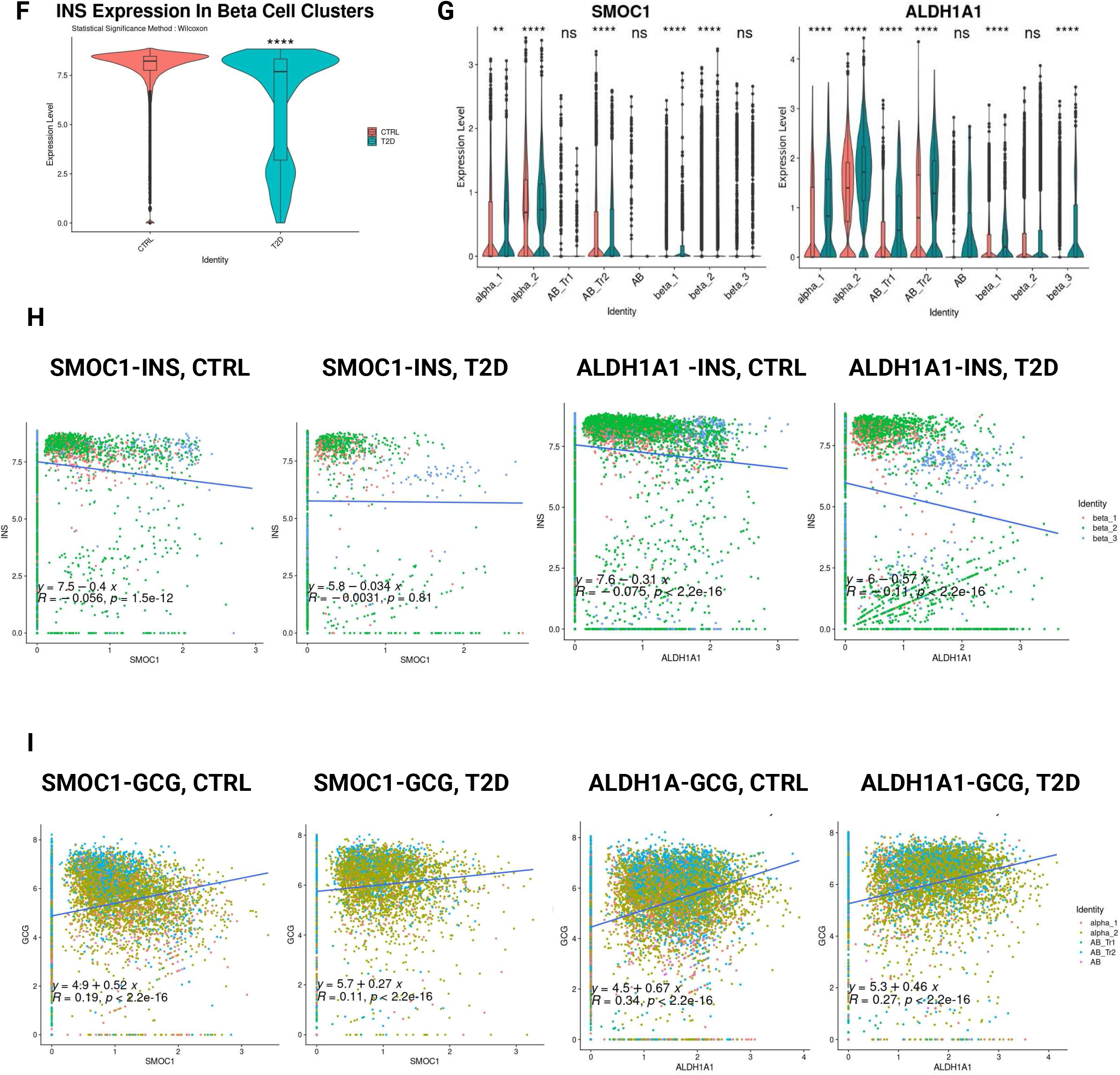
Velocity Analysis, Gene Expression and Gene Correlation in HPAP/HIRN Non-Diabetic and T2D Human Islet Dataset. **A.** Splicing ratio of α-and β-subclusters in control (left) and in T2D data (right). **B.** Scale matched velocity length of non-diabetic (left) and T2D human islet cell data (right). **C.** Velocity stream by scVelo analysis. UMAP coordinate and subcluster annotation was obtained by projecting data to our scRNA/snRNA-seq reference. **D.** PAGA trajectory with stochastic modeling for non-diabetic (left) and T2D islet data (right) **E.** *INS* expression in β-cells in non-diabetic and T2D islet data and analyzed statistically by Wilcoxon rank-sum test. **F.** Expression of *SMOC1* and *ALDH1A1* in α-and β-cell subclusters, split by diabetic condition (red for non-diabetic and green for T2D islets). **G.** Correlation of *SMOC1* and *ALDH1A1* expression with *INS* expression in β-cell subclusters. **H.** Correlation of *SMOC1* and *ALDH1A1* with *GCG* in α-cell subclusters.

RNA velocity stream and PAGA analysis showed that non-diabetic human α- and β-cells exhibited a relatively complex multi-directional pattern with trajectories in both directions from α- to β-cells and vice versa, implying mutual plasticity among α- and β-cell subclusters (**Fig. 7C-E**), in part resembling the pattern observed in human islet samples *in vitro* with scRNA- and snRNA-seq analysis (**Fig. 3B-E**). In contrast to non-diabetic human islets, human T2D α- and β-cells followed a clear unidirectional β- to α-cell transition, moving from β1 to β3, then to AB and finally to α2 cells or direct transition from β1 to α1 and then to α2 (**Fig. 7C-D**). This is exactly opposite of the transition observed *in vivo* in human islet grafts and suggests pressure on β-cells to become dedifferentiated or converted to α-cells in subjects with T2D.

### β-to-α cell trajectory genes and T2D

T2D is characterized by a decrease in β-cells potentially due to increased apoptosis and dedifferentiation (39–41). In the HIRN-HPAP database, we found that β-cells in T2D islets display decreased *INS* expression (**Fig. 7E**). We then explored whether any of the common genes involved in β-to-α cell transition (**Fig. 4F**) might be upregulated in β-cells in T2D islets directing their phenotype towards α-cells. Among these genes, *ZNF311*, *PAPPA2, TM4SF4* and *PLCE1* showed little or no increase in their expression in β-cells in T2D islets (**Fig. 7F and Suppl. Fig. 4A**). Of the remaining genes, *NR4A1, PCSK2, SLC30A8, and BTG2* expression showed positive correlation with *INS* expression in T2D β-cells (**Suppl. Fig. 4B**). All the genes showed a positive correlation between their expression and *GCG* expression in α-cells (**Suppl. Fig. 4C**). Finally, two genes *SMOC1* and *ALDH1A1* were upregulated in β-cells of T2D islets and negatively correlated with *INS* expression in non-diabetic β-cells while positively correlated with *GCG* in α-cells (**Fig. 7G-I, Suppl. Fig. 4C**).

## DISCUSSION

The use of single cell and single nucleus transcriptomics can help to define specific α-cell and β-cell subtypes involved in the development and progression of T2D (2,8,12–17). It can also uncover genes that can participate in the β-cell dedifferentiation or the β- to α-cell transition processes that occur in T2D providing additional therapeutic targets for diabetes treatment. Furthermore, deciphering specific genes and pathways in α-cell subpopulations that can enable their reprograming into insulin-producing cells for β-cell replacement therapies can also facilitate the identification of new targets for diabetes treatment (1–6,8,18–22).

We have recently reported that three different human β-cell subpopulations with their corresponding distinct transcriptome profiles can be identified using integrated scRNA-seq and snRNA-seq analysis of human islets (17). The proportion of these human β-cell subpopulations changes significantly depending on whether the analysis is done in human islets *in vitro* or in human islet grafts *in vivo* (17). Extending this earlier dataset from human islets *in vitro*, we have now identified five different *GCG*-expressing α-cell subclusters - α1, α2, AB-Tr1, AB-Tr2, and AB - displaying different transcriptome profiles. α2 cells are enriched in mitotic/cell cycle pathways suggesting that these might be the cycling α-cells recently described (42–44). The *GCG*-expressing AB subcluster is a multihormonal gene expression cluster enriched in genes related to pancreatic progenitor and β-cell pathways while α1, α2, AB-Tr1, and AB-Tr2 clusters are enriched in genes specific for α-cell functions and translation pathways. RNA velocity and PAGA analysis identify AB cells as the root of the transition to both α- and β-cells at least in isolated human islets. Using single cell transcriptomics and pseudotime analysis, Saikia et al have identified four human α-cell subclusters with a varying degree of *GCG* expression, two of them displaying mature gene cell markers, one with β-cell like features (*INS* and *GCG* expression) and one intermediate (45). Recently, a transcriptional cross species map of pancreatic islet cells from mouse, pig and humans has annotated four α-cell states in which 50% were mature α-cells and almost 20% had an immature or precursor-like profile (46). Our findings in human islets *in vitro* resemble these previous studies with similar proportions of mature (α1+α2 cells) and precursor-like profile cells (AB-Tr1+AB cells). However, these previous studies were performed only in isolated islets, and they did not address whether these α-cell subtypes exist *in vivo* (45,46).

To our surprise, we find that AB cells are not present *in vivo* in human islet grafts where 96% of α-cells are α1 plus α2 cells, the mature α-cells. *In vivo*, α2 cells are not enriched in mitotic/cell cycle pathways suggesting these cells lose the identity of cycling α-cells in this setting (42–44). Our results also suggest that AB cells are a product of islet isolation and *in vitro* culture conditions while *in vivo* most of the α-cells in human islet grafts are mature α-cells with a small proportion of cells with progenitor capacity, similar to results we previously observed with human β-cell subpopulations (17). This suggests that *in vivo*, at least in the human islet graft environment, the number of α- and β-cells in human islets with progenitor potential is limited. It is important to note that the multihormonal cells at a transcriptional level exist *in vivo* in human islet grafts (α-cell subtypes AB-Tr1 and AB-Tr2) but highly reduced in their proportion (4%). We speculate that this limited number of cells could represent the small percentage of α-cells that can be reprogrammed to β-cells with acute β-cell loss, as observed in mice (18). In contrast to isolated islets *in vitro*, trajectory inference analysis of α- and β-cells in human islet grafts favors only unidirectional α-to-β-cell transition. This is not surprising since, at least in mice, acute β-cell loss leads to transdifferentiation of α-cells into β-cells (18) while acute α-cell loss does not initiate a transdifferentiation program from β-cells to α-cells (38). Our studies in human islet grafts with snRNA-seq analysis align with potential transcriptional transition from human α-cells to β-cells but not the opposite, at least under basal conditions.

Gene commonality analysis of the trajectories from AB to α-cells (trajectories B, C and D) identifies *ZNF331, PLCE1, PAPPA2, TM4SF4, NR4A1, BTG2, SLC30A8, PSCK2, SMOC1,* and *ALDH1A1* as exclusive common genes in the three trajectories. *ZNF331* encodes for zinc finger protein 331 and has been identified as a putative tumor suppressor silenced or downregulated in gastrointestinal cancers (47), but its function in islet cells remains unknown. Phospholipase C-ɛ-1 (PLCE1) links Epac2 activation to the potentiation of glucose-stimulated insulin secretion (GSIS) in mouse islets (48) and its expression is decreased in β-cells while maintained in α-cells in T2D islets (49). In our dataset, in T2D islets, we find that *PLCE1* expression increases in α1 and α2 (mature α-cells) and decreases further from the already very low expression level in mature β1 cells confirming previous results and highlighting this phospholipase as a potential candidate for β-cell dysfunction in T2D. Pappalysin-2 (PAPPA2), an enzyme that cleaves and inactivates IGFBP proteins, is upregulated in α-cells during gestation (50). In our study, we observe that *PAPPA2* is expressed in α-cells and may participate in the transition of AB/β-cells into α-cells by suppressing the effects of IGFBPs since IGFBP1 has been reported to promote α- to β-cell transdifferentiation (51). Thus, PAPPA2 could suppress the effect of IGFBPs favoring the α-cell phenotype. We also observe that *PAPPA2* is upregulated in α-cells of T2D islets suggesting that PAPPA2 upregulation can limit the activity and potential beneficial effects of IGFBPs. Interestingly, a prospective human study has shown that high IGFBP1 levels reduce the risk of developing T2D by more than 85% (51,52) perhaps reflecting PAPPA2 upregulation in T2D. Transmembrane 4 L Six Family Member 4 (TM4SF4) is a tetraspanin protein localized at the membrane of human α-cells but not β-cells (53). Loss-of-function studies have revealed that tm4sf4 loss inhibits α and β cell specification (54). In the current studies, we observe that *TM4SF4* is upregulated in α-cells in T2D islets suggesting further specification of endocrine cells into α-cells. However, its function in adult α-cells is unknown. Although we observed a relative increase of *TM4SF4* expression in β-cells, its expression is still very low, and the biological importance of this statistically significant increase is unknown. The orphan nuclear receptor NR4A1 is critical for fuel utilization and mitochondrial function in liver, muscle, and adipose tissues and is important for β-cell mitochondrial function and insulin secretion (55). However, *NR4A1* expression is not altered in islets from T2D subjects (56). This is in contrast with the observation in our single cell transcriptomics study in which *NR4A1* expression is increased in α-cells compared with β-cells in non-diabetic islets and it is upregulated in β-cells of T2D subjects, perhaps as a compensatory mechanism for insulin resistance in T2D. B-cell translocation gene 2 (BTG2) is upregulated by high glucose levels and induces apoptosis in pancreatic β-cells (57). However, its role in α-cells is unknown. Here we observe that *BTG2* expression is unaltered in α-cells but enhanced in β-cells of T2D islets perhaps suggesting involvement in the β-cell death process that occurs in T2D (58). *SLC30A8* encodes the Solute Carrier Family 30 (Zinc Transporter), Member 8, an islet zinc transporter restricted to α- and β-cells which is responsible for zinc accumulation into secretory granules (59). Importantly, loss-of-function mutations in *SLC30A8* in humans protect against T2D, suggesting SLC30A8 inhibition as a therapeutic strategy in T2D prevention (60). Indeed, in our studies we observe that *SLC30A8* expression is present in both mature α- and β-cells and increased in T2D β-cells. *PCSK2* encodes the pre-pro-glucagon processing enzyme. Its expression is higher in α-cells than β-cells in our studies, as expected, and is increased in T2D islet cells. It has been reported that T2D β-cells have elevated PCSK2 immunoreactivity, apparently not contributing to impaired proinsulin processing (61). Therefore, *PCSK2* appears to be required for transition from β-cells to α-cells but its role in human β-cells is not completely known. Collectively, this novel gene set merits further exploration in α- and β-cells in human and animal models of T2D-associated islet dysfunction.

Two additional common genes in the trajectories from β-to-α-cells are the aldehyde dehydrogenase 1 isoform A1 (*ALDH1A1*) and the SPARC Related Modular Calcium Binding 1 (*SMOC1*). In general, Aldh isoform expression and enzymatic activity mark progenitor populations in adult and embryonic mouse pancreas and have been identified as a potent regulator of pancreatic endocrine differentiation (62,63). *ALDH1A1* is mostly expressed in α-cells as shown in a previous report (53) and in our studies. The isoform ALDH1A3 has been implicated in β-cell dedifferentiation in T2D (39–41), however the involvement of the α-cell isoform ALDH1A1 in β-cell dedifferentiation in T2D has not previously been reported. We observe that the expression of *ALDH1A1* increases in all the β-cell subtypes in T2D and that it negatively correlates with *INS* expression. These results clearly position ALDH1A1 as a mediator of, or participant in β-to α-cell conversion and suggest that this family of enzymes may serve as therapeutic targets for diabetes treatment (41). SMOC1 has been previously identified as a secreted protein regulator that affects islet function in obese T2D mouse islets (64). In our dataset, *SMOC1* is mostly expressed in α-cells in basal conditions. Its expression in β-cells negatively correlates with *INS* expression, and it is enhanced in mature β-cells in T2D suggesting a potential role in β-cell dedifferentiation. Collectively these studies highlight the power of common gene analysis in trajectory inference in single cell transcriptomics studies to identify genes that could participate in T2D development and could be therapeutic targets for treating the disease.

Along these lines, gene commonality analysis of the trajectories from AB to β-cells also identified *ZNF385D, TRPM3, CASR, HDAC9* and *MEG3* as common genes in these trajectories. TRPM3 (Transient Receptor Potential Melastatin 3) channels are non-selective cation channels expressed in insulinoma cells and pancreatic β-cells that may participate in GSIS (65). In our studies, *TRPM3* expression correlates with the GSEA pathway of pancreatic β-cell development which confirms its potential involvement in AB to β-cell transition. However, only a small percentage of cells express this gene, and the detection is mostly observed by snRNA-seq suggesting its pre-mRNA nature. The Calcium-Sensing Receptor (CASR) plays an important role in cell-to-cell communication in pancreatic islet cells by detecting local concentrations of extracellular Ca2+ in the intra-islet space and enhancing GSIS (66,67). As with *TRPM3*, *CASR* expression correlates with the GSEA pathway of β-cell development, but in this case, it is highly expressed in ∼50% β1 and β2 cells in scRNA-seq suggesting the presence of mature RNA in these cells. The long non-coding (lnc) RNA-maternal expressed gene 3 (*MEG3*) is an imprinted gene expressed in β-cells and downregulated in T1D and T2D (68). Knockdown of *Meg3* in mouse β-cells impairs insulin synthesis and secretion, enhances apoptosis, and decreases Pdx-1 and MafA mRNA and protein expression, suggesting a role as regulator of β-cell identity (68). In humans, the Kaestner group has identified an imprinted region on chromosome 14, the *DLK1-MEG3* locus, as being downregulated in islets from T2DM (69). Collectively, these reports and our results suggest that *MEG3* lncRNA may have important roles in maintaining human β-cell identity and function and may be essential for the transition of progenitor cells (AB cells) to β-cells, a subject not studied thus far. In addition to these three β-cell function and identity genes, we found that zinc finger 385D (*ZNF385*) is also a common gene in the trajectory towards β-cells and its expression correlates with enhanced β-cell differentiation by GSEA. *ZNF385* is one of the top expressed genes that defines β-cells in snRNAseq, as we have previously reported (17). Importantly, Gene-Wide Association Studies have strongly associated *ZNF385D* gene variants with T2D onset (70,71) and forced expression of *ZNF385D* together with retinoic acid receptor β (*RARB*) and Growth Arrest Specific 7 (*GAS7*) genes in non-diabetic human islets leads to the appearance of a T2D-β-like phenotype in these islets (72). Interestingly, *ZNF385D* expression occurs mostly in the snRNA-seq dataset (pre-mRNA). Thus, while its pre-mRNA presence can be indicative of a β-cell phenotype, its splicing and mature mRNA expression can result in dedifferentiated β-cells (72), illustrating the importance of discerning differences driven by gene expression, splicing and/or function. This suggests that studying RNA splicing combined with snRNA-seq analysis can be of great importance to determine specific gene functions in islet cell subpopulations. Histone Deacetylase 9 (HDAC9) is one of the class-II HDACs and a transcriptional co-repressor, which is involved in neuronal development and pancreatic β cell differentiation (73). Genetic deletion of *HDAC9* increases β-cell numbers at embryonic and neonatal ages in mice while having no effect on α-cell mass (74,75). It is unknown at this point whether this occurs in adult mice as well. Interestingly, in adult non-diabetic human islets, *HDAC9* is mainly expressed in β-cells and at lower levels in α-cells and it has been suggested that β-cell-specific HDAC9 expression in the pancreas could be a reprogramming agent and enforcer of β-cell identity (76). Another set of common genes in the transition from AB to β-cells includes the obligated *INS* and *IAPP* genes. Interestingly, the *SST* gene is also part of this common set of genes, but its expression is restricted to β3 cells, the most immature β-cells perhaps suggesting a bifurcation from β3 towards mature β-cells (β1 and β2) or δ-cells, an aspect that warrants further studies. Finally, retinol-binding protein 4 (RBP4), a plasma retinol transporter from hepatocytes to peripheral tissues, is a diabetogenic adipokine elevated in plasma of patients with T2D (77). Although the role of RBP4 in the human β-cell remains elusive, the potential importance of this retinol binding protein in pancreatic β-cell dysfunction in rodents has been suggested (78). Interestingly, a recent study comparing gene expression in human and mouse α- and β-cells using scRNA-seq has shown that *RBP4* expression is very low in human α-cells but robust in mouse α-cells with an opposite expression pattern in human and mouse β-cells (79). These current studies clearly indicate differences in the pattern of expression depending on the species, confirm RBP4 as a human β-cell gene, and support the need for studies in human islets to define the role of this retinol-binding protein in β-cells.

In summary, these studies illustrate the importance of this integrated scRNA-seq and snRNA-seq reference to distinguish different α-cell subtypes and their potential transcriptional status. These studies also provide evidence of the potential transcriptional transition among α-cell and β-cell subtypes depending on the pathophysiological context. They also highlight the utility of these omics platforms for finding potential gene targets such as *SMOC1* and *ALDH1A1* which might modulate β-cell dedifferentiation in T2D and could be therapeutic targets for treating the disease. Finally, and perhaps most importantly, they illustrate the importance of studying human islet genomics and transcriptomics in settings as close as possible to their native *in vivo* environment, as compared to artificial culture systems and following damaging human islet isolation and individual islet cell dispersion commonly used for scRNA-seq studies. The human islet *in vivo* graft model used here, and recently developed human pancreatic slice model (80) both coupled with snRNAseq, provide ideal next-generation platforms for transcriptomic analysis.

## ACKNOWLEDGEMENTS

We thank the NIDDK-supported Einstein-Sinai DRC and the Human Adenovirus and Islet Core for help with the proposed studies and Prodo Labs for supplying human cadaveric islets. We acknowledge support from NIH grants P30 DK020541, R01 DK126450, R01 DK116873, R01 DK116904, R01 DK125285, R01 DK105015, R01 DK129196, R01 DK130300, K-01 DK128378, and an Einstein-Sinai DRC Pilot and Feasibility Grant (to GL).

## SUPPLEMENTAL FIGURE LEGENDS

**Supplemental Figure 1: α-Cell Subpopulations, Differentially Expressed Genes and Pathways. A.** Number of cells in each α-cell subpopulation. **B.** Differentially expressed genes for each α-cell subpopulation; α1 with *NEAT1, ACTG1, PEAK1, ACTB*; α2 with *FAP, PCSK2, SLC30A8, GLS;* AB_Tr1 with *HSPA1A, HSPH1, DNAJB1, PLCG2*; AB_Tr2 with *PCSK1N, GAPDH, TTR, SNHG29*; AB with *INS, SST, IAPP, PPY*. **C.** Differentially enriched pathways for each subpopulation.

**Supplemental Figure 2: Differential Pathway Enrichment Analysis for Human Islet Grafts *In Vivo* snRNA-seq α-Cell subclusters.**

**Supplemental Figure 3: Trajectory Associated Gene Expression along the Pseudotime in the α-β-Cell Transition**. We selected 100 genes with high pseudo-time association, clustered by the expression pattern.

**Supplemental Figure 4: Correlation between Expression of Common Genes and *INS* and *GCG*. A.** Expression of common genes between non-diabetic (red) and T2D (green) islet data and analyzed statistically by Wilcoxon rank-sum test. **B.** Correlation of expression of common genes in the α- to β-cell trajectories with *INS* expression in β-cell subclusters. **C.** Correlation of expression of common genes in the α- to β-cell trajectories with *GCG* in α-cell subclusters.

