## Supplementary material for "Human Pancreatic α-Cell Heterogeneity and Trajectory Inference Analysis Using Integrated Single Cell- and Single Nucleus-RNA Sequencing Platforms": Table 1

### Supplemental Table 1:

### Checklist for Reporting Human Islet Preparations Used in Research

Adapted from Hart NJ, Powers AC (2018) Progress, challenges, and suggestions for using human islets to understand islet biology and human diabetes. Diabetologia https://doi.org/10.1007/s00125-018-4772-2

| **Islet preparation** | **1** | **2** | **3** | **4** | **5** | **6** | **7** |
|---|---|---|---|---|---|---|---|
| Unique identifier | HP20240 | HP20245 | HP21189 | HP20234 | HP21055 | HP21203 | HP21197 |
| Donor age (years) | 42 | 24 | 26 | 58 | 36 | 25 | 53 |
| Donor sex (M/F) | M | M | M | F | F | M | M |
| Donor BMI (kg/m$^2$) | 23.5 | 20.0 | 26.3 | 26.9 | 31.6 | 26.5 | 32.4 |
| Donor $HbA_{1c}$ or other measure of blood glucose control | 5.4% | 5.4% | 5.4% | 5.5% | 5.6% | 5.8% | 5.5% |
| Origin/source of islets[b] | Prodo Lab | Prodo Lab | Prodo Lab | Prodo Lab | Prodo Lab | Prodo Lab | Prodo Lab |
| Islet isolation centre | Prodo Aliso Viejo, CA | Prodo Aliso Viejo, CA | Prodo Aliso Viejo, CA | Prodo Aliso Viejo, CA | Prodo Aliso Viejo, CA | Prodo Aliso Viejo, CA | Prodo Aliso Viejo, CA |
| Donor history of diabetes? Please select yes/no from drop down list | No | No | No | No | No | No | No |
| Diabetes duration (years) | | | | | | | |
| Glucose-lowering therapy at time of death[c] | | | | | | | |

*Continues on the next page*

| | | | | | | | |
|---|---|---|---|---|---|---|---|
| Donor cause of death | Stroke | Head trauma | Head trauma | Stroke | Stroke | Head trauma | Anoxic event |
| Warm ischaemia time (h) | N/A | N/A | N/A | N/A | N/A | N/A | N/A |
| Cold ischaemia time (h) | N/A | N/A | N/A | N/A | N/A | N/A | N/A |
| Estimated purity (%) | 95 | 95 | 90 | 90 | 90 | 95 | 90 |
| Estimated viability (%) | 95 | 95 | 95 | 95 | 95 | 95 | 95 |
| Total culture time (h)[d] | 120 | 120 | 120 | 96 | 96 | 144 | 120 |
| Glucose-stimulated insulin secretion or other functional measurement[e] | N/A | N/A | N/A | N/A | N/A | N/A | N/A |
| Handpicked to purity? Please select yes/no from drop down list | Yes | Yes | Yes | Yes | Yes | Yes | Yes |
| Additional notes | | | | | | | |

[a]If you have used more than eight islet preparations, please complete additional forms as necessary
[b]For example, IIDP, ECIT, Alberta IsletCore
[c]Please specify the therapy/therapies
[d]Time of islet culture at the isolation centre, during shipment and at the receiving laboratory
[e]Please specify the test and the results

| **Islet preparation** | **8** | **9** | **10** | **11** |
|---|---|---|---|---|
| Unique identifier | HP21280 | HP21292 | HP21302 | HP22214 |
| Donor age (years) | 69 | 44 | 72 | 46 |
| Donor sex (M/F) | Male | Female | Male | Male |
| Donor BMI (kg/m$^2$) | 24.54 | 32.2 | 20.4 | 23.1 |
| Donor $HbA_{1c}$ or other measure of blood glucose control | 5.8 | 5.7 | 5.1 | 4.9 |
| Origin/source of islets[b] | Prodo Lab | Prodo Lab | Prodo Lab | Prodo Lab |
| Islet isolation centre | Prodo Aliso Viejo, CA | Prodo Aliso Viejo, CA | Prodo Aliso Viejo, CA | Prodo Aliso Viejo, CA |
| Donor history of diabetes? Please select yes/no from drop down list | No | No | No | No |
| Diabetes duration (years) | | | | |
| Glucose-lowering therapy at time of death[c] | | | | |

*Continues on the next page*

| | | | | |
|---|---|---|---|---|
| Donor cause of death | Head taruma | Stroke | Anoxic Event | Stroke |
| Warm ischaemia time (h) | N/A | N/A | N/A | N/A |
| Cold ischaemia time (h) | N/A | N/A | N/A | N/A |
| Estimated purity (%) | 90 | 85 | 85 | 90 |
| Estimated viability (%) | 95 | 95 | 95 | 95 |
| Total culture time (h)[d] | 144 | 144 | 144 | 144 |
| Glucose-stimulated insulin secretion or other functional measurement[e] | N/A | N/A | N/A | N/A |
| Handpicked to purity? Please select yes/no from drop down list | Yes | Yes | Yes | Yes |
| Additional notes | | | | |
