## Supplemental Figures for "Human Pancreatic α-Cell Heterogeneity and Trajectory Inference Analysis Using Integrated Single Cell- and Single Nucleus-RNA Sequencing Platforms"

### Supplemental Figure 1

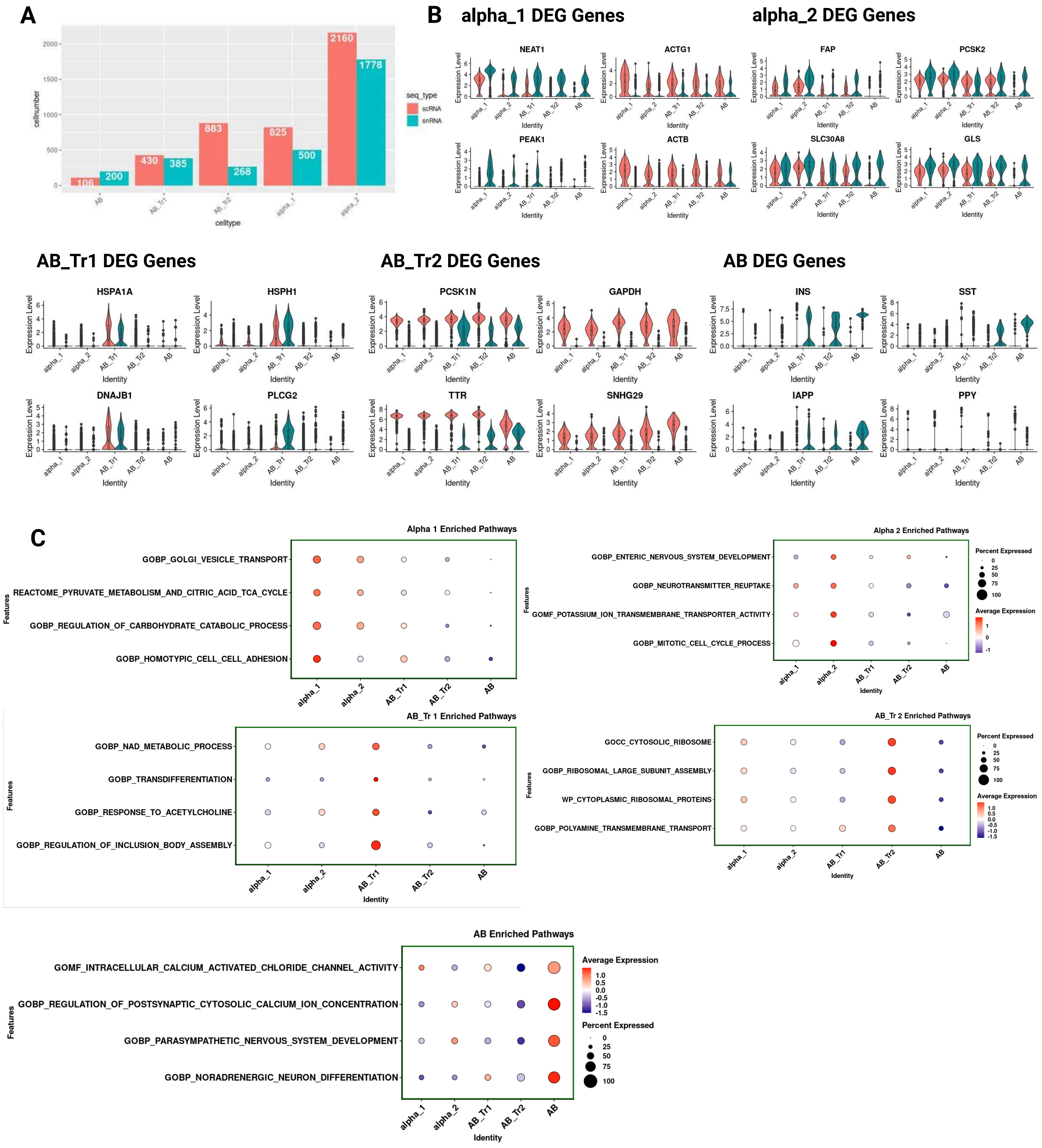

### Supplemental Figure 2

A

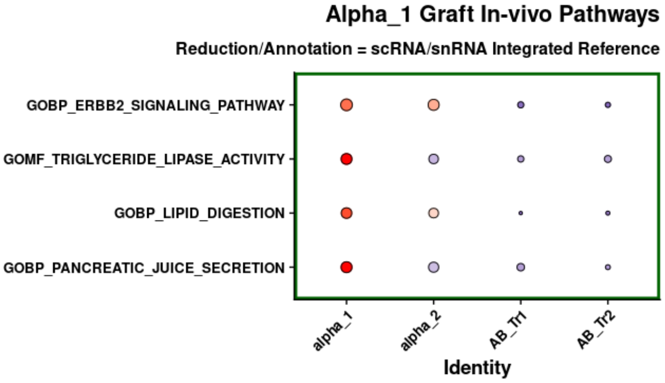


B

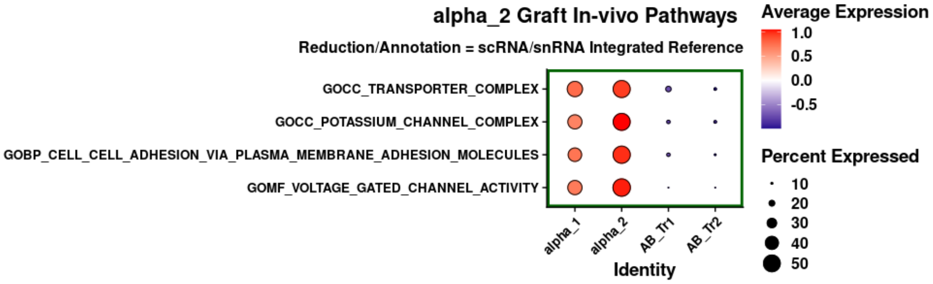


C

D

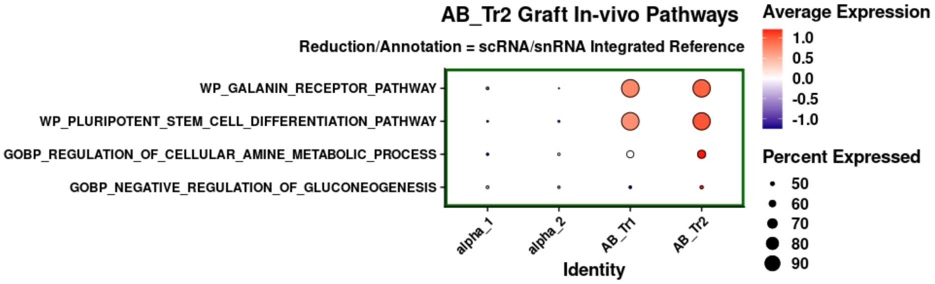

### Supplemental Figure 3

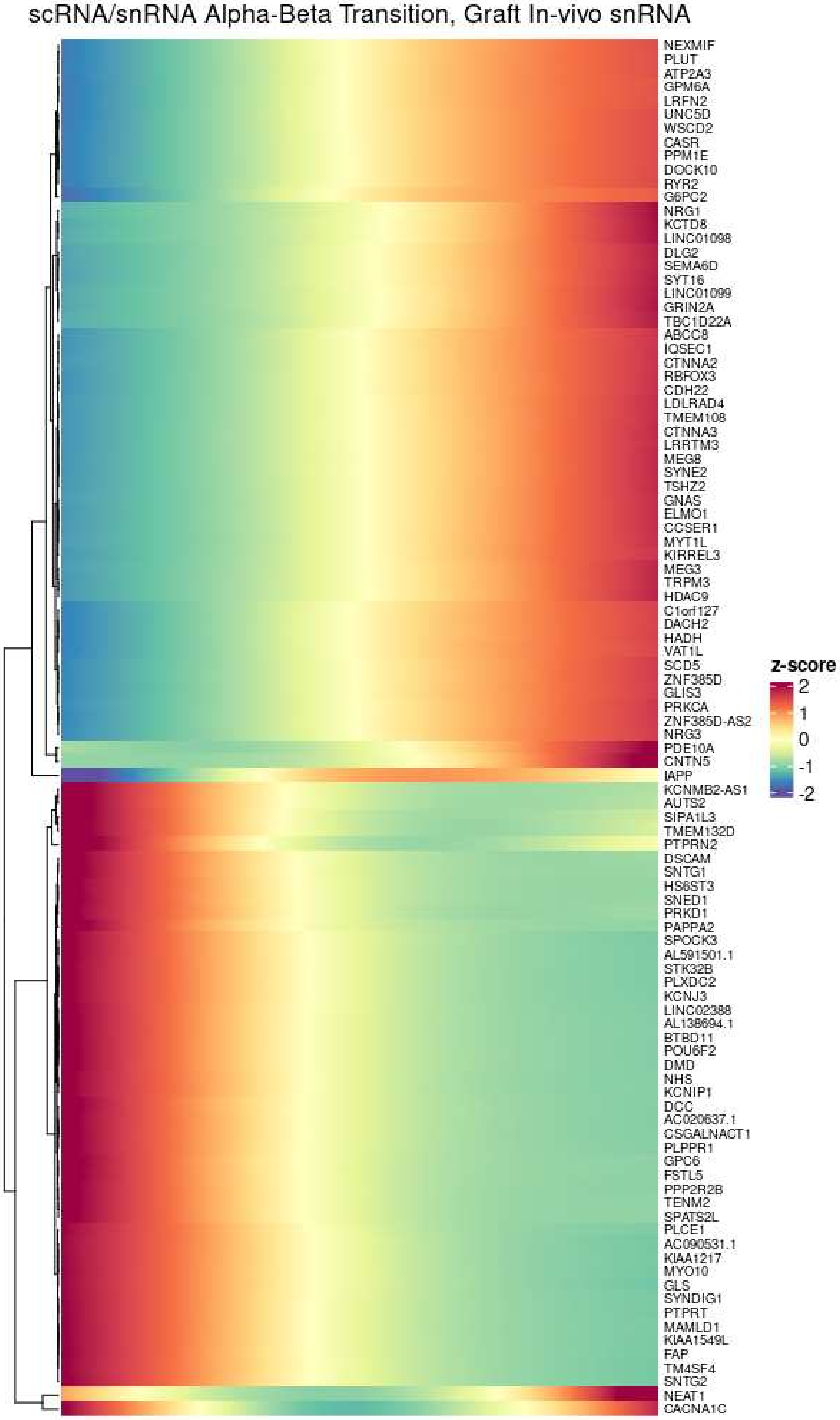

### Supplemental Figure 4

A

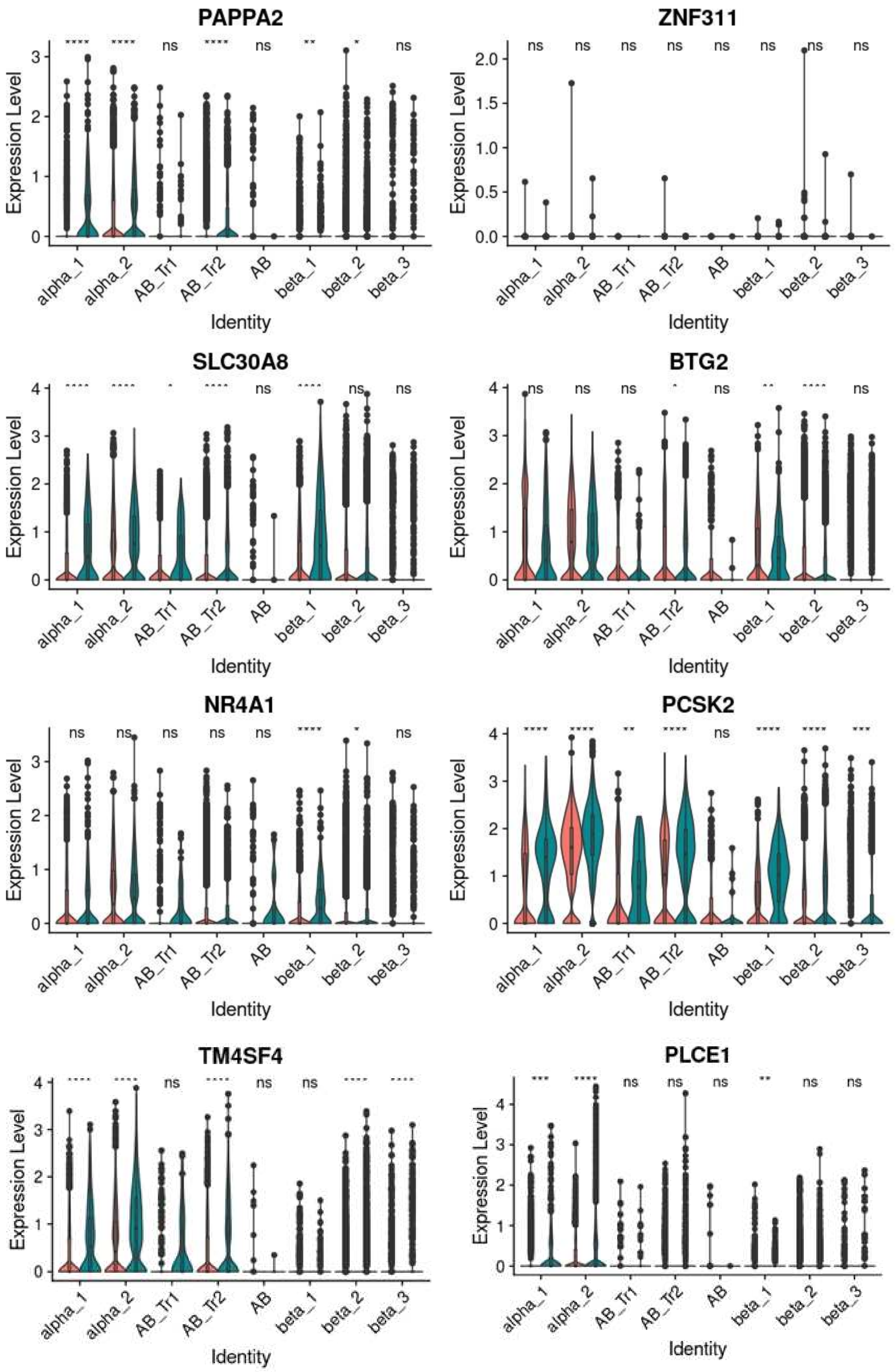

### Supplemental Figure 4 (Continuation)

## B

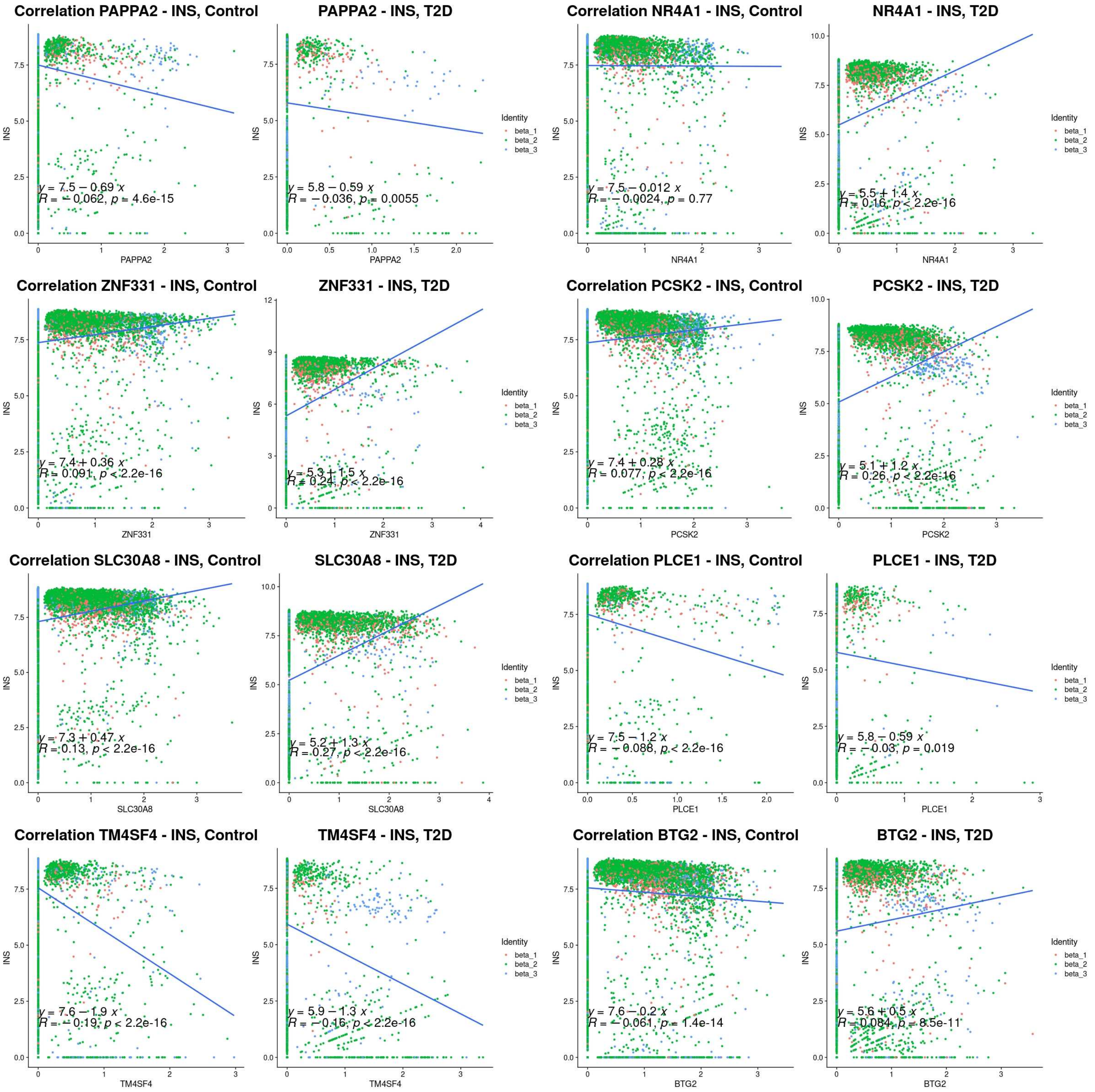

### Supplemental Figure 4 (Continuation)

**C**

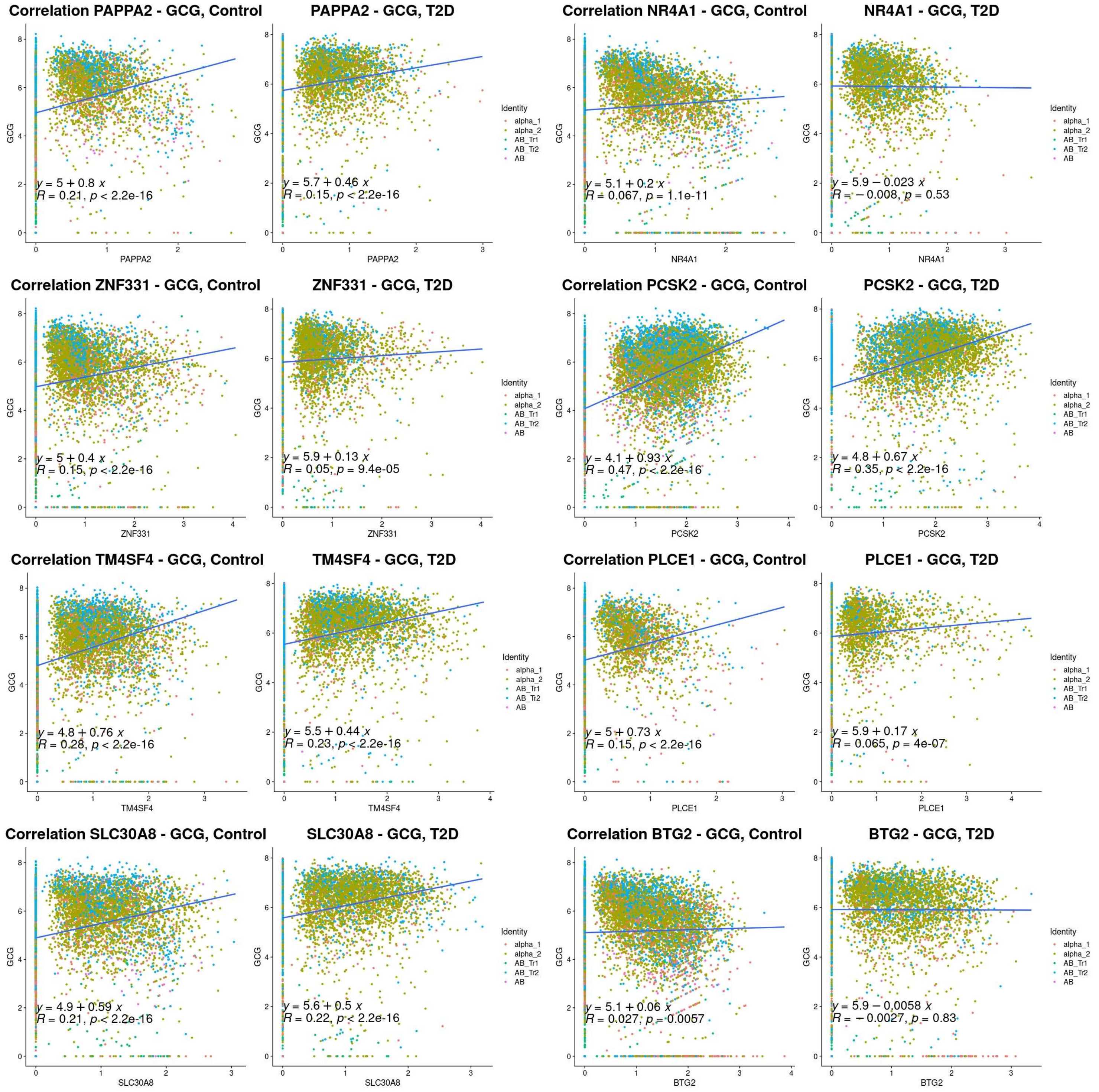
